# Pancreatic cancer cells assemble a CXCL12-keratin 19 coating to resist immunotherapy

**DOI:** 10.1101/776419

**Authors:** Zhikai Wang, Ran Yan, Jiayun Li, Ya Gao, Philip Moresco, Min Yao, Jaclyn F. Hechtman, Matthew J. Weiss, Tobias Janowitz, Douglas T. Fearon

## Abstract

How pancreatic ductal adenocarcinoma (PDA) cells stimulate CXCR4 to exclude T cells and resist T cell checkpoint inhibitors is not known. Here, we find that CXCL12, the ligand for CXCR4 that is produced by the cancer-associated fibroblast, “coats” human PDA and colorectal cancer cells as covalent heterodimers with keratin 19 (KRT19). Modeling the formation of the heterodimer with three proteins shows that KRT19 binds CXCL12 and transglutaminase-2 (TGM2), and that TGM2 converts the reversible KRT19-CXCL12 complex into a covalent heterodimer. We validate this model by showing that cancer cells in mouse PDA tumors must express KRT19 and TGM2 to become coated with CXCL12, exclude T cells, and resist immunotherapy with anti-PD-1 antibody. Thus, PDA cells have a cell-autonomous means by which they capture CXCL12 to mediate immune suppression, which is potentially amenable to therapy.

**One Sentence Summary:** Cancer cells in pancreatic ductal adenocarcinoma use transglutaminase-2 to assemble a coating comprised of covalent CXCL12-keratin 19 heterodimers that excludes T cells and mediates resistance to inhibition of the PD-1 T cell checkpoint.

## Main Text

The major challenge in cancer immunology is to define the molecular pathway by which adenocarcinomas exclude T cells and resist immunotherapy with inhibitors of T cell checkpoints (*1*). An early clue to the identity of this pathway was the finding that depleting the cancer-associated fibroblast (CAF) in an autochthonous mouse model of PDA slowed tumor growth and uncovered sensitivity to the anti-tumor effects of anti-PD-L1 antibody (*2*). The immune suppressive function of the CAF was traced to its production of the chemokine, CXCL12, when treatment of PDA-bearing mice with AMD3100, a small molecule inhibitor of CXCR4, the receptor for CXCL12, overcame the exclusion of T cells by cancer cell nests, and uncovered the anti-tumor activity of anti-PD-L1 antibody. The capacity of CXCR4 that has been stimulated by CXCL12 to suppress the intra-tumoral accumulation of T cells may be related to its inhibition of the chemotactic function of CXCR3 (*3*). Two recent early stage clinical studies of CXCR4 inhibition in patients with metastatic PDA have confirmed the clinical relevance of these pre-clinical observations (*3, 4*).

Since CXCR4 signaling accounts for the non-random distribution of intra-tumoral T cells away from cancer cells and into the stroma, we determined whether CXCL12 also was present in a non-random, but opposite, distribution in human PDA and other carcinomas. We stained sections of freshly resected human PDA and colorectal cancer (CRC) with fluorochrome-conjugated antibodies to KRT19, which would identify cancer cells, and to CXCL12. Confocal microscopy showed that only the KRT19^+^ cancer cells demonstrated binding of the anti-CXCL12 antibody (Fig. 1A). This restricted distribution of CXCL12 to cancer cells was observed in seven additional types of human carcinomas (fig. S1A and B). Human melanoma, which is a non-epithelial cancer that does not express KRT19, did not bind anti-CXCL12 antibody.

**Figure 1:**
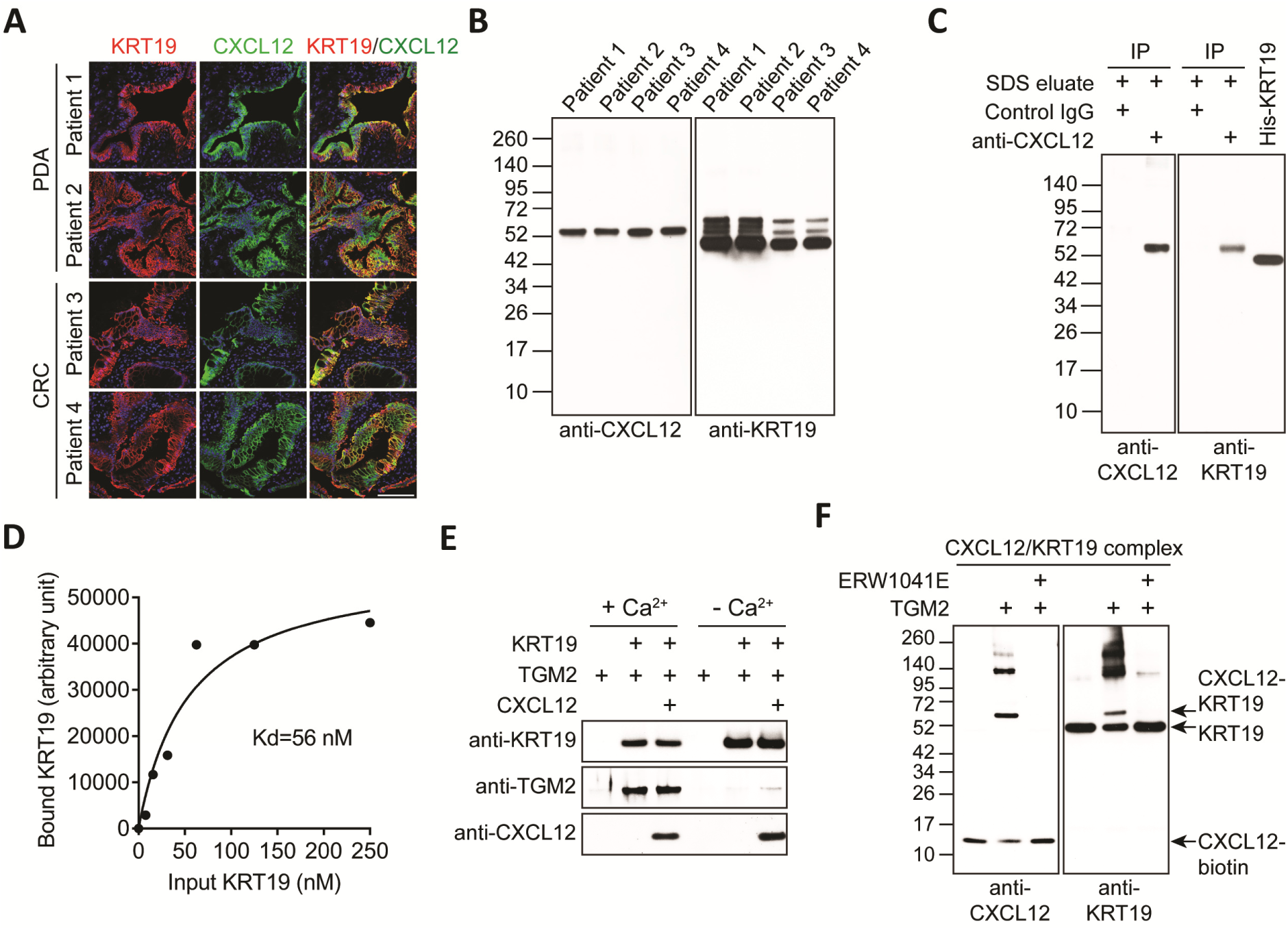
The CXCL12-KRT19 coating of cancer cells. (**A**) Sections of freshly resected human PDA and CRC were stained with fluorochrome-conjugated antibodies to KRT19 and CXCL12. Scale bar, 100 µm. (**B**) SDS-eluates of samples of the resected PDA and CRC were 5 subjected to SDS-PAGE and immunoblotting with antibodies to CXCL12 and KRT19. (**C**) The proteins that were immunoprecipitated by control IgG or anti-CXCL12 IgG from the pooled SDS-eluates of the four tumors were subjected to SDS-PAGE and immunoblotting with anti-CXCL12 and anti-KRT19 antibodies. (**D**) Streptavidin-beads bearing CXCL12-biotin were incubated with increasing concentrations of KRT19. Bound KRT19 was detected by SDS-PAGE 10 and immunoblotting with anti-KRT19 antibody. The apparent affinity with which CXCL12 and KRT19 interacted was calculated. (**E**) Anti-KRT19 antibody/protein G beads were incubated at 4°C with combinations of KRT19, CXCL12 and TGM2, in the presence or absence of Ca^2+^. The bound proteins were detected by SDS-PAGE and immunoblotting with the indicated antibodies. (**F**) Streptavidin beads bearing preformed complexes of biotinylated CXCL12 and KRT19 were 15 incubated at 20°C for 15 min with TGM2 in the presence or absence of the TGM2 inhibitor, ERW1041E. The proteins were eluted from the beads and detected by SDS-PAGE and immunoblotting with antibodies to CXCL12 and KRT19.

To determine how CXCL12 associates with cancer cells in PDA and CRC, we sequentially eluted samples of the human PDA and CRC tumors with buffers containing NP-40 and SDS, respectively, and analyzed the eluted proteins by SDS-PAGE followed by immunoblotting with anti-CXCL12 and anti-KRT19 antibodies. CXCL12, despite being a soluble protein that is secreted by CAFs, was detected only in the SDS-eluates (Fig. 1B and fig. S1C). Moreover, the eluted CXCL12 appeared with a relative molecular weight of ∼55 kD instead of the nominal 10-13 kD molecular weights of its isoforms. KRT19, as expected for an intermediate filament protein, also required SDS for solubilization, and presented with its predicted molecular weight of ∼45 kD, and two higher molecular weight forms, one of which corresponded to the molecular weight of the eluted CXCL12 (Fig. 1B). Examination of the immunoprecipitate obtained with anti-CXCL12 antibody from the combined SDS-eluates of the four tumors revealed that this high molecular weight form of KRT19 co-immunoprecipitated with anti-CXCL12, indicating the occurrence of a CXCL12-KRT19 heterodimer (Fig. 1C). Mouse PDA, in which KRT19^+^ cancer cells also stain with fluorochrome-conjugated anti-CXCL12 antibody (fig. S2A), also demonstrated the presence of an SDS-soluble, CXCL12-KRT19 heterodimer (fig. S2B and C). Thus, all CXCL12 that could be detected in the SDS-eluates of human PDA and CRC and mouse PDA was present in a stable, apparently covalent, heterodimer with KRT19.

Confirming a previous finding (*2*), *Cxcl12* mRNA was detected in mouse PDA tumors mainly in CAFs expressing *Pdgfra* mRNA (fig. S2D). Individual cancer cells in non-permeabilized sections could be stained with antibodies specific for CXCL12 and KRT19, respectively (fig S2E). Also, analysis by flow cytometry of intact single cells from mouse PDA tumors that had been stained with anti-CXCL12 and anti-KRT19 antibodies showed that all CXCL12^+^ cells were KRT19^+^ (fig. S2F). Thus, cancer cells are capable of secreting the intermediate filament protein, KRT19, as has been previously reported (*5*), to participate in the formation of a CXCL12-KRT19 heterodimer.

We modeled the formation of the CXCL12-KRT19 complex by reasoning that a non-covalent interaction between KRT19 and CXCL12 is stabilized by an enzyme capable of crosslinking proteins, such as TGM2, which is expressed by both PDA and CRC cancer cells (*6, 7*). Streptavidin beads bearing C-terminally biotinylated human CXCL12 were incubated with increasing concentrations of KRT19, and bound KRT19 was measured by SDS-PAGE and immunoblotting with anti-KRT19 antibody. CXCL12 bound KRT19 in a saturable manner with a nanomolar affinity (Fig. 1D and fig. S3A). As a negative control, KRT19 did not interact with another chemokine, CXCL8 (fig. S3B). Formation of a trimolecular complex was then assessed by incubating beads bearing KRT19 at 4°C with CXCL12 and TGM2 in the presence or absence of Ca^2+^. KRT19 bound CXCL12 regardless of the presence of Ca^2+^, but binding of TGM2 required the cation, suggesting that the open conformation of TGM2 was required for this interaction (Fig. 1E). Beads bearing biotinylated CXCL12 only bound TGM2 in the presence of KRT19 and Ca^2+^, confirming the Ca^2+^-dependence of the interaction between KRT19 and TGM2 (fig. S3C). Preformed CXCL12-KRT19 complexes were incubated with TGM2 at 20°C in the presence of Ca^2+^, and with or without ERW1041E, an inhibitor of the TGM2 transamidase site (*8*). TGM2 generated a heterodimeric complex of ∼55kD containing CXCL12 and KRT19, similar to complex that was eluted from PDA and CRC, and its formation was blocked by ERW1041E (Fig. 1F). An additional, higher molecular form of CXCL12 and KRT19 was also generated that was not observed in the SDS-eluates of tumors, perhaps indicating that that SDS does not solubilize all CXCL12-containing complexes from tumors. Including ERW1041E during the detergent elution of mouse PDA tumors did not diminish recovery of the high molecular weight form of CXCL12, excluding a possibility that the complex was generated *in vitro* (fig. S3D). Thus, KRT19 has two independent sites that mediate the binding of CXCL12 and of TGM2 in its Ca^2+^-induced, catalytically active open conformation (*9*). This trimolecular interaction may facilitate the formation of the CXCL12-KRT19 coating of cancer cells.

To test the prediction that TGM2 is required for the formation of the CXCL12-KRT19 coating *in vivo*, we generated subcutaneous tumors with control PDA cells or PDA cells in which the *Tgm2* gene had been edited (fig. S4), in wild type mice or mice in which the *Tgm2* gene had been interrupted (*10*). Analysis of tumor sections revealed that loss of TGM2 from cancer cells, but not from the host, greatly diminished both the staining of cancer cells with anti-CXCL12 antibody (Fig. 2A), and the amount of the SDS-soluble, high molecular weight form of CXCL12 (fig. S5A). The residual high molecular weight CXCL12 may have been generated by FXIIIa, a transglutaminase that is expressed by macrophages and was detected by immunofluorescence (fig. S5B), but not by other members of the transglutaminase family, which are not expressed by mouse PDA cells (fig.S5C) (*11*). T cells accumulated in areas of the tumors in which CXCL12 was absent, and there was T cell-dependent slowing of tumor growth that was accompanied by elevated mRNA levels of genes characteristic of cytotoxic CD8^+^ T cells (Fig. 2B-D). Despite the presence of host-derived TGM2 in the stroma of PDA tumors (fig. S5D), this TGM2 did not compensate for loss of cancer cell TGM2 in the coating of cancer cells with CXCL12, nor affect tumor growth (Fig. 2C). Thus, TGM2 from the cancer cells has an essential role in formation of the CXCL12-KRT19 coating.

**Figure 2:**
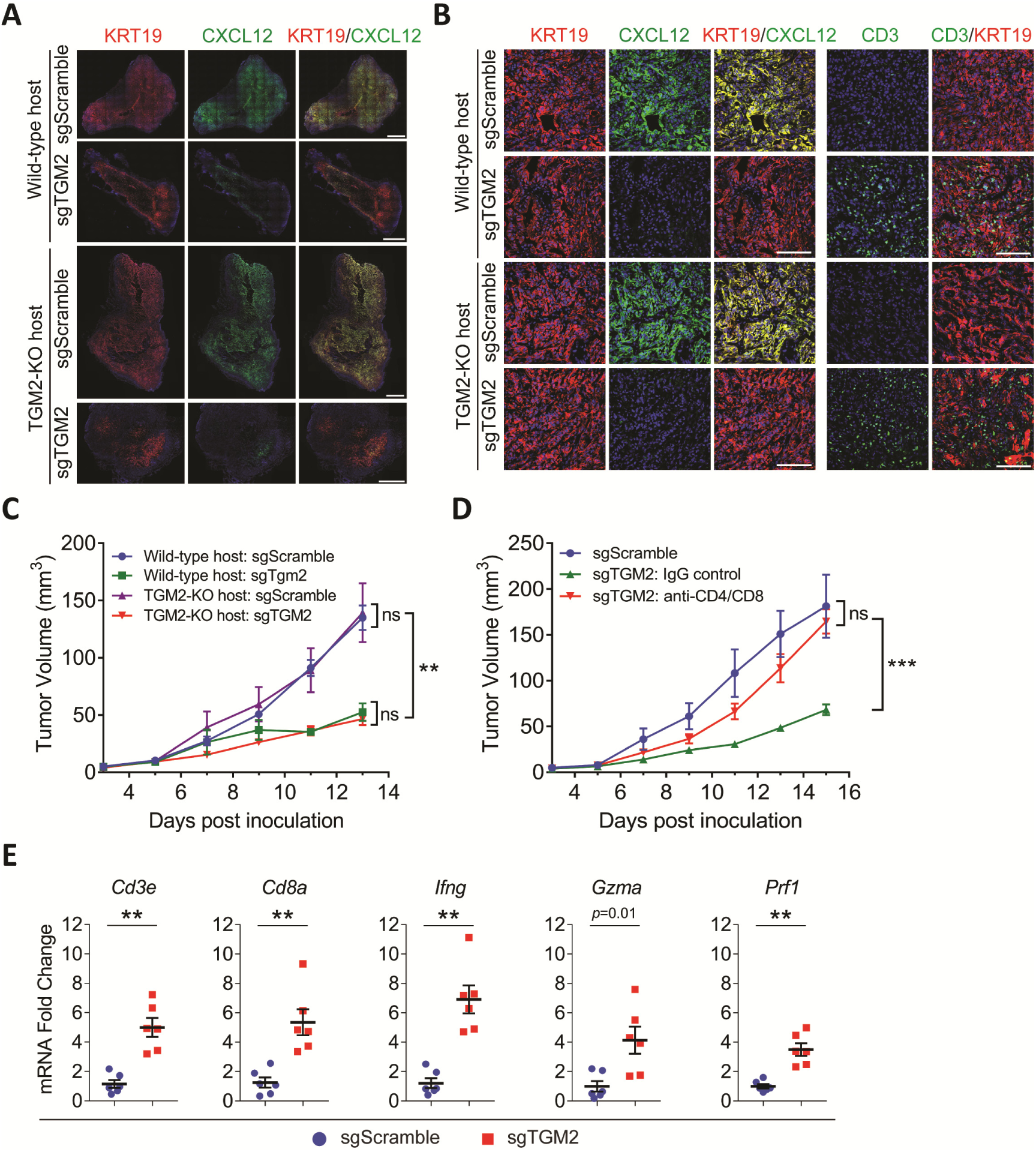
The role of TGM2 expressed by PDA cells in the formation of the CXCL12-KRT19 coating. sgScramble or sgTGM2-edited PDA cells were inoculated subcutaneously in wild-type hosts or TGM2-deficient hosts. (**A**) Cross sections of the entire tumors of each type were stained with fluorochrome-conjugated antibodies to KRT19 and CXCL12. Representative images are shown. Scale bars, 1 mm. (**B**) Sections of each of the four tumor types were stained with fluorochrome-conjugated antibodies to KRT19, CXCL12, and CD3. Representative images are shown. Scale bars, 100 µm. (**C**) The growth curves of each of the four tumor types in panel B are shown. n=5. (**D**) The growth curves are shown of tumors formed with sgScramble PDA cells and sgTGM2-edited PDA cells, respectively, in wild-type hosts that were untreated, or treated with non-immune IgG or the T cell depleting antibodies to CD4 and CD8. n=5. (**E**) The mRNA levels of immune genes in tumors formed with sgTGM2-edited PDA cells in wild-type hosts were compared to those of tumors formed with sgScramble PDA cells. Mean ± SEM; ns, not significant, ** *p*<0.01, *** *p*<0.001, Student’s *t* test.

An indispensable role for KRT19 was demonstrated by the absence of the CXCL12-KRT19 coating on PDA cells in which the *Krt19* gene had been edited (Fig. 3A and fig. S6A-C). These tumors also were infiltrated with T cells, exhibited T cell-dependent, slower growth, and had elevated mRNA levels of immunologically relevant genes (Fig. 3A-C). In contrast, interrupting the expression of KRT18 in PDA cells did not diminish the staining of tumors with anti-CXCL12 antibody (fig. S6D). Rescue of KRT19 expression in the PDA cells reversed these phenotypic changes (fig. S7). Therefore, we conclude that the CXCL12-KRT19 heterodimer mediates T cell exclusion and immune suppression, since blocking its formation by interrupting the expression of two unrelated genes, *Tgm2* and *Krt19*, results in the same immunological outcome.

**Figure 3:**
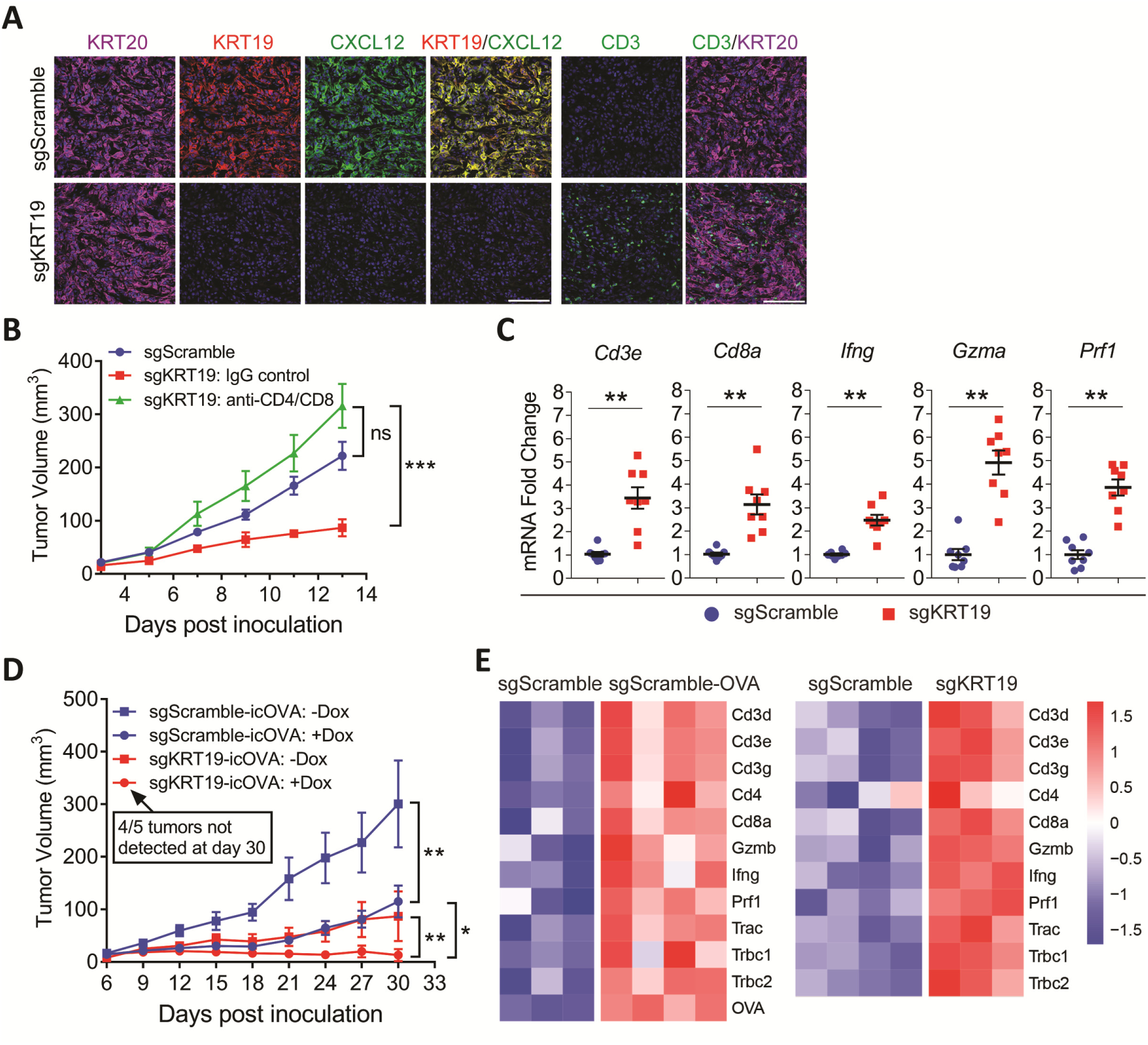
The role of KRT19 expressed by PDA cells in the formation of the CXCL12-KRT19 coating. (**A**) Sections of subcutaneous tumors formed with sgScramble control PDA cells and sgKRT19-edited PDA cells were stained with fluorochrome-conjugated antibodies to KRT19, KRT20, CXCL12 and CD3. Scale bars, 100 µm. (**B**) The growth curves are shown of subcutaneous tumors formed with sgScramble PDA cells and sgKRT19-edited PDA cells in mice that were untreated or treated with either non-immune IgG or T cell-depleting antibodies to CD4 and CD8. n=5. **(C)** The mRNA levels of immune genes in tumors formed with sgKRT19-edited PDA cells are compared to those of tumors formed with sgScramble PDA cells. (**D**) The growth curves are shown of subcutaneous tumors formed with sgScramble PDA cells or sgKRT19-edited PDA cells that had been engineered to express doxycycline (Dox)-inducible cytoplasmic OVA (icOVA) in mice fed normal food (-Dox) or food containing Dox (+Dox). Mean ± SEM; n=5. ns, not significant, *, *p*<0.05, ** *p*<0.01, *** *p*<0.001. Paired *t* test. (**E**) The heat map is shown of the relative levels of expression of immune genes as detected by RNA-Seq assay of individual subcutaneous tumors formed with sgScramble PDA cells, OVA-expressing sgScramble PDA cells, and sgKRT19-edited PDA cells, respectively.

T cells accumulate in cancer cell nests in microsatellite instable (MSI) PDA (*12*), yet the CXCL12-KRT19 coating is maintained (fig. S8A, B), suggesting either that this cancer represents an exception to the immune suppressive function of the CXCL12-KRT19 coating, or that the increased immunogenicity of MSI PDA is an independent determinant of the intra-tumoral immune reaction. To distinguish between these possibilities, we formed hepatic metastases and subcutaneous tumors containing mouse PDA cells expressing the foreign antigen, ovalbumin (OVA). The OVA-expressing hepatic metastases were infiltrated with OVA-specific CD8^+^ T cells (fig. S8C), reproducing this characteristic of MSI PDA. In subcutaneous tumors, expression of OVA or loss of KRT19 equivalently slowed tumor growth, and led to the increased expression of the same set of immune-related genes (Fig. 3D and E). Combining OVA expression with the absence of KRT19 expression further decreased PDA growth so that 4/5 tumors were eliminated by day 30. Thus, immune suppression by the CXCL12-KRT19 coating is maintained in a highly immunogenic PDA tumor, and immunogenicity and the CXCL12-KRT19 coating are independent determinants of the intra-tumoral immune reaction.

A role for the CXCL12-KRT19 coating in the resistance of mouse PDA to inhibition of the PD-1 T cell checkpoint was assessed. Subcutaneous tumors were formed with control PDA cells or cells that did not express either KRT19 or TGM2. Treatment with rat anti-PD-1 IgG or non-immune IgG was initiated on day 13 when all PDA tumors were established. The administration of these rat antibodies was limited to one week, after which mouse anti-rat IgG antibodies develop (fig. S9). The growth rate of tumors containing control PDA cells was not altered by the administration of the anti-PD-1 antibody, and the experiment was terminated on day 17 when tumors had attained a pre-determined size limit (Fig. 4A). Tumors comprised of PDA cells lacking expression of either KRT19 or TGM2 grew more slowly than those formed with control PDA cells, and responded to anti-PD-1 antibody with diminished growth rates (Fig. 4A) and increased transcription of genes expressed by effector CD8^+^ T cells (fig. S10). Hepatic metastases formed with luciferase-expressing control PDA cells exhibited staining with anti-CXCL12 antibody and excluded T cells, whereas those formed with KRT19-deficient PDA cells lacked the CXCL12-KRT19 coating and were infiltrated with T cells, despite containing CXCL12-expressing CAFs (fig. S11). Treatment with anti-PD-1 antibody did not alter the growth rates of KRT19-expressing tumors, whereas it arrested the growth of metastases containing PDA cells not expressing KRT19 (Fig. 4B and fig. S12), and increased the expression of genes indicating the presence of effector CD8^+^ T cells (fig. S13).

**Figure 4:**
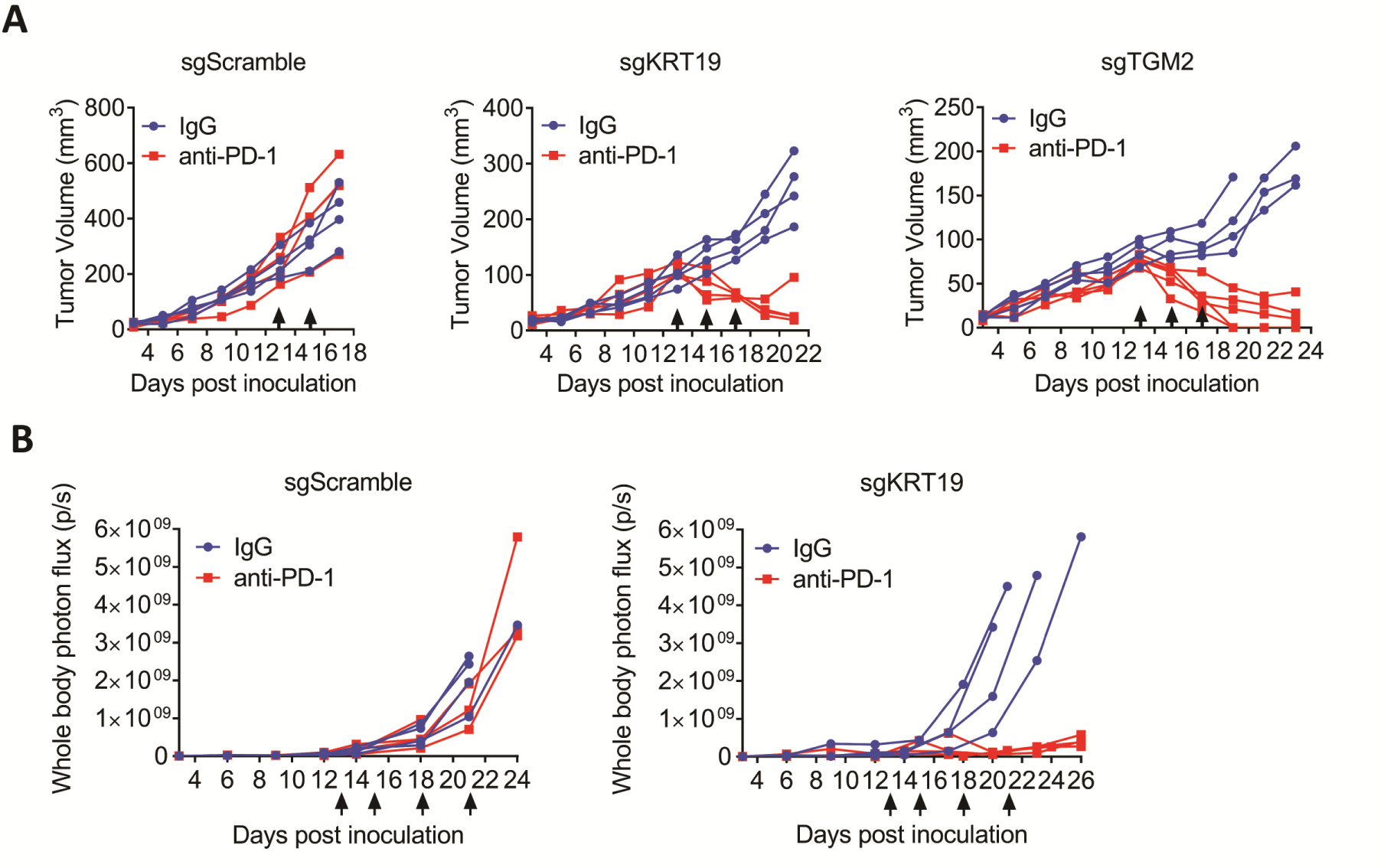
Treatment of mice bearing PDA tumors with antibody to PD-1. (**A**) Mice bearing subcutaneous tumors formed with sgScramble control PDA cells, sgKRT19-edited PDA cells, or sgTGM2-edited PDA cells were treated by intra-peritoneal administration of non-immune IgG or anti-PD-1 IgG (arrows), and tumor growth was measured. (**B**) Mice bearing hepatic metastases formed with luciferase-expressing sgScramble control PDA cells or sgKRT19-edited PDA cells were treated with non-immune IgG or anti-PD-1 IgG (arrows), and growth of the metastases was measured by bioluminescent imaging.

Taken together, these studies show that there is a cancer cell-autonomous mechanism involving KRT19 and TGM2 for capturing CXCL12, the mediator of immune suppression by CAFs. The resulting CXCL12-KRT19 coating explains the capacity of cancer cells to exclude T cells and to resist inhibitors of the PD-1 checkpoint. The finding that KRT19 interacts with CXCL12 and TGM2 to assemble the CXCL12-KRT19 coating offers new approaches to cancer immunotherapy.

## Acknowledgments

We thank Dr. David Tuveson for providing the 1242 and mM1 PDA cells. We thank Bruno Gegenhuber for help with RNAseq analysis. We also acknowledge the contributions of the Cold Spring Harbor Laboratory facilities of Animal, Microscopy, Flow Cytometry, and Animal Imaging.

## Author contributions

D.T.F. and Z.W. conceived the project. Z.W., R.Y., J.L., Y.G., P.M., M.Y., J.F.H., and M.J.W. performed the experiments; D.T.F., T.J., Z.W., R.Y., J.L., Y.G. and P.M. analyzed data; D.T.F., and Z.W. wrote the manuscript; D.T.F., T.J., Z.W., R.Y., J.L., Y.G., P.M. and M.Y. revised the manuscript.

## Data and materials availability

All data are available in the manuscript or the supplementary material.

## Supplementary Materials

### Materials and Methods

#### Human Tumor Samples

All human tumor samples used in this study are approved by institutions’ Institutional Review Board (IRB). Freshly resected human pancreas and colon tumors were obtained from Northwell Health Tissue Donation Program. The collected tumor samples were temporarily stored in RPMI medium and delivered on ice the same day of surgical removal. After receiving, part of each sample was embedded in Tissue-Tek OCT compound (Sakura, 4583) for immunofluorescence staining and the rest was flash-frozen in liquid nitrogen for protein extraction. Formalin fixed paraffin embedded (FFPE) human tumor arrays for pancreas cancer (PA501a), colon cancer (BC05021a), breast cancer (BC08013c), bladder cancer (BC12011d), melanoma (ME483a) and frozen multiple organ tumor/tissue arrays (FMC282e and FMO401) were purchased from US Biomax.

#### Tumor Protein Extraction

Tumor pieces of about 100 mg were used for protein extraction. Freshly dissected tumor pieces or the tumor pieces that were previously flash-frozen in liquid nitrogen and subsequently stored at −80°C were briefly homogenized in 1 ml ice cold IP buffer (75 mM HEPES, pH 7.5, 150 mM NaCl, 1 mM DTT, 10% Glycerol and protease inhibitor cocktail (Thermo, 78437)) plus 1% NP-40. After removal of remaining large tissue particles by flowing through the Pierce Tissue Strainer (Thermo, 87791), the homogenized tissue was sequentially extracted using the following step-wise instructions: (1) Re-suspend the tissue and incubate at 4°C on rotator for 10 min followed by centrifugation at 4,000 g for 5 min. Save the supernatant as “NP-40 extract”. Repeat to wash the pellet; (2) Re-suspend the pellet in 250 μl IP buffer plus 0.1% sodium deoxycholate (DOC) and incubate at 4°C on rotator for 30 min followed by centrifugation at 6,000 g for 5 min. Save the supernatant as “DOC extract”. Repeat to wash the pellet; (3) Re-suspend the pellet in 200 μl IP buffer plus 5 mM CaCl_2_ and 3,000 units/ml micrococcal nuclease (Thermo, 88216) and incubate at room temperature for 30 min followed by centrifugation at 16,000 g for 5 min. Save the supernatant as “DNase extract”. Repeat to wash the pellet; (4) Re-suspend the pellet in 150 μl IP buffer plus 1% SDS and incubate at room temperature for 10 min followed by centrifugation at 16,000 g for 5 min. Keep the supernatant as “SDS extract”. This is the fraction of cytoskeleton. For the extractions with TGM2 inhibition, 100 μg/ml or 500 μg/ml ERW1041E (Sigma, 509522) or equivalent volume of DMSO was included in every extraction buffer. Protein concentration was measured using a protein assay kit (BioRad, 5000112) and 1 μg of each fraction were boiled in Laemmli sample buffer containing 20 mM DTT, and applied to SDS-PAGE gel for western blot.

#### Immunoprecipitation of CXCL12-KRT19 coating

SDS in the cytoskeleton fraction was removed with the Pierce Detergent Removal Spin Column (Thermo, 87777). For immunoprecipitation, 100 μg of the protein extract was diluted in 500 μl IP buffer containing 1% BSA (Calbiochem, 2930) and 0.1% SDS which helps to prevent self-assembly of cytokeratin proteins. The mixture was pre-cleared by centrifugation at 16,000 g for 10 min and the supernatant was transferred in to a new tube. Then, 5 μg Rabbit isotype IgG (Thermo, 10500C) or Rabbit anti-CXCL12 antibody (LSBio, LS-C48888) was added in to the solution and incubated on rotor at 4°C overnight. Separately, 100 ul protein G agarose beads (Cell signaling, 37478, 50% slurry) were pre-blocked in IP buffer containing 1% BSA. After overnight incubation, 20 μl pure protein G resins (or 40 μl 50% slurry) were added directly to the protein/antibody solution and incubated for another 2 hours at 4°C. Following three washes with IP buffer containing 0.1% SDS, the resins were boiled in 100 μl Laemmli sample buffer containing 20 mM DTT, and 10 μl of each sample was applied to SDS-PAGE gel for western blot where Veriblot for IP detection reagent HRP (Abcam, ab131366) was used.

#### Protein Binding Assays and Cross-linking

The buffer of commercially available recombinant human KRT19 protein (Creative Biomart, KRT19-7239H) was changed to protein binding buffer (25 mM Tris, pH 7.5, 100 mM NaCl, 1% Triton X-100), using Zeba spin desalting columns (Thermo, 89882). For protein binding, biotinylated recombinant human CXCL12 (Chemotactics, B-CXCL12) or CXCL8 (Chemotactics, B-CXCL8) was incubated with KRT19 at 4°C for 2 hours. Then, streptavidin conjugated Dynabeads (Thermo, 11205D), pre-blocked with 1% BSA, were added and incubated for another 2 hours at 4°C. Following three washes, bound proteins were boiled in Laemmli sample buffer containing 20 mM DTT for western blotting. To measure the binding affinity between CXCL12 and KRT19, 100 ng CXCL12-biotin bound on streptavidin Dynabeads was incubated with 0, 8 nM, 16 nM, 31 nM, 63 nM, 125 nM or 250 nM KRT19 at 4°C for 4 hours. For TGM2 binding, KRT19 or KRT19/CXCL12 (R&D Systems, 350-NS) complex was firstly immobilized on protein G Dynabeads (Thermo, 10004D) via anti-KRT19 antibody (Abcam, ab76539). Alternatively, CXCL12-biotin or KRT19/CXCL12-biotin complex was immobilized on streptavidin Dynabeads. Then, recombinant human TGM2 (R&D Systems, 4376-TG-050) was added and incubated in the presence or absence of 10 mM CaCl_2_ on ice for 10 minutes.

Following one or two quick washes on ice, with calcium containing binding buffer, bound proteins were boiled for western blotting.

To do KRT19-CXCL12 cross-linking, 100 ng CXCL12-biotin and 500 ng KRT19 was incubated and bound on streptavidin Dynabeads. After three washes, KRT19/CXCL12-biotin complex bound Dynabeads were re-suspended in 50 μl TGM2 activation buffer (100 mM Tris, pH 7.5, 150 mM NaCl, 10 mM CaCl_2,_ 10 mM DTT). Then, 50 ng recombinant human TGM2, together with or without 100 μM ERW1041E, was added and incubated at room temperature for 15 minutes. The reactions were stopped by direct boiling in Laemmli sample buffer containing 20 mM DTT.

#### Animals

C57BL/6 mice of 6 weeks age were purchased from The Jackson Laboratory. After receiving, the mice were housed in a helicobacter free room for 2 weeks before each experiment. CD45.1 OT-I mice were generated by cross-breeding OT-I mice (Jackson, 003831) and B6.SJL mice (Jackson, 002014). TGM2-KO mice were obtained by cross-breeding *Tgm2*-floxed mice (Jackson, 024694) to transgenic CMV-Cre mice (Jackson, 006054). The progeny were genotyped at Transnetyx for double-allele knockouts. All animal experiments were approved by the Cold Spring Harbor Laboratory (CSHL) Institutional Animal Care and Use Committee (IACUC) in concordance with the NIH “Guide for the Care and Use of Laboratory Animals”.

#### CRISPR Editing and Lentivirus

The control CRISPR guide (sgScramble: GCTTAGTTACGCGTGGACGA (*13*)) and the guides targeted to mouse *Krt19* gene (sgKRT19-1: CACAGGAAATTACTGCCCTG; sgKRT19-2: TAGTGGTTGTAATCTCGGGA; sgKRT19-3: CGGAGGACGAGGTCACGAAG) or mouse *Tgm2* gene (sgTGM2-1: GAATATGTCCTTACGCAACA; sgTGM2-2: GTCCTGTTGGTCCAGCACTG; sgTGM2-3: TTGACCTCGGCAAACACGAA) were cloned into the lentiCRISPR-v2 plasmid (Addgene, 52961) and sequencing validated. The lentivirus was produced by co-transfecting HEK293T cells with the guide containing lentiCRISPR-v2 plasmid together with the lentivirus packing plasmid psPAX2 (Addgene, 12260) and the envelop expressing plasmid pMD2.G (Addgene, 12259).

#### Cancer Cell Lines

Mouse pancreas ductal adenocarcinoma (PDA) cancer cell lines 1242 and mM1 which were derived from KPC (LSL-Kras^G12D/+^; LSL-Trp53^R172H/+^; Pdx1-Cre) mouse tumor (*14*) were a gift from the laboratory of Dr. Tuveson. To generate sgScramble, sgKRT19 and sgTGM2 cell lines, 1242 cells were transduced with lentivirus expressing SpCas9 and specific CRISPR guides. The transduced cells were directly selected with 1 μg/ml puromycin (Sigma, P8833) for polyclonal stable cell lines. The single cell clones were generated by limited dilution. To rescue KRT19 expression in the *Krt19*-edited cancer cells which was achieved by transduction with the lentivirus containing the sgKRT19-1 guide, the single cell clone was further transduced with lentivirus expressing mouse KRT19 with four synonymous mutations at the guide targeting site. To facilitate tumor imaging *in vivo* for hepatic metastasis, the sgScramble and sgKRT19 cell lines were transduced with lentiviral plasmid expressing firefly luciferase and GFP which are linked by 2A peptide. The luciferase/GFP transduced cells were sorted by FACS. Doxycycline-inducible ovalbumin-expressing cells were generated by transducing mM1cells, 1242-sgScramble cells and 1242-sgKRT19 cells with the lentiviral vector pCW57-gfp-p2a-mcs (Addgene, 89181) containing the cytoplasmic version of ovalbumin followed by FACS sorting and single cell cloning. All the cancer cell lines were cultured in DMEM medium (Cellgro, 10-013-CV) supplemented with 10% FBS (Seradigm, 1500-500), 100 units/ml penicillin and 100 μg/ml streptomycin.

#### Mouse Studies

For subcutaneous tumors, 2.5×10^5^ cells, unless indicated otherwise, were injected into the right flank of mice. Tumor volume was measured with caliper every two or three days and calculated according to the formula V = (L×W^2^)/2. Mice that were developing tumor associated skin ulcer during the experiment were humanely sacrificed, and the visually ulcerated tumors were not included in further analysis. For the hepatic metastasis model, 2.5×10^4^ or 1.0×10^5^ luciferase expressing cells were injected into the portal vein. Tumor size was monitored through bioluminescence imaging every 3 or 4 days. To do so, 150 μl of 30 mg/ml D-Luciferin (Perkin-Elmer, 122799-10) was administered through intraperitoneal injection and mice were imaged with the IVIS Spectrum *in vivo* Imaging System. For mouse tumor immunotherapy, 200 μg isotype IgG (BioXcell, BP0089) or Rat anti-mouse PD-1 antibody (BioXcell, BP0273) was given through intraperitoneal injection on indicated days. T cell depletion was achieved by intraperitoneal injection of 200 μg anti-CD4 (BioXcell, BP0003) and anti-CD8 (BioXcell, BP0061) antibodies twice a week. Mice that were developing tumor associated ascites or lethargy during the experiment were humanely sacrificed.

As to the ovalbumin inducible hepatic metastasis model, 1000 CD45.1 OT-I CD8^+^ cells, isolated from the spleen of CD45.1 OT-I mouse using CD8a^+^ T cell isolation kit (Miltenyi Biotec, 130-104-075), were injected into C57BL/6J mice intravenously prior to cancer cell injection via the portal vein. Doxycycline induction of ovalbumin was 11 days after cancer cell injection (5×10^4^ cells), using doxycycline food, and the metastatic tumors were harvest 7 days later. For the ovalbumin inducible 1242-sgScramble and 1242-sgKRT19 subcutaneous tumors, 5×10^5^ cells were inoculated and ovalbumin was induced at day 0.

#### Immunofluorescence

Mouse tumors were embedded in Tissue-Tek OCT compound (Sakura, 4583) and freshly frozen on dry ice. In the case of hepatic metastatic tumors, the whole livers were perfused through the portal vein with cold PBS before freezing. The embedded tumors were sectioned at 10 μm for staining. For Human FFPE tumor arrays, the slides were deparaffinized in Xylene for three times (5 min each) followed by sequential rehydration treatments in 100% ethanol (5 min, 3 times), 95% ethanol (10 min, twice), 70% ethanol (10 min, twice), 50% ethanol (10 min, twice), and water (5 min, twice). After rehydration, antigen retrieval was conducted by boiling the sample in 10 mM Tris-HCl (pH 8.8) plus 1 mM EDTA for 10 min, followed by cooling down to room temperature for 30 min and subsequent two washes in water for 5 min each.

To stain tumor sections, the samples were fixed with freshly prepared PLP buffer (100 mM phosphate buffer, pH 7.4, 80 mM L-lysine, 10 mM NaIO_4_, and 4% paraformaldehyde) for 10 min at room temperature. After washing with PBS, a hydrophobic barrier was drawn around the tumor slice with ImmEdge PAP pen (Vector Laboratories, H-4000). The sections were then permeabilized with 0.2% TritonX-100 in PBS for 15 min followed by blocking with 1% BSA (Calbiochem, 2930) plus 10% normal goat serum (Thermo, 16210064) in PBS at room temperature for 1 hour. After the blocking, the tumor slices were stained by incubation with non-conjugated primary antibodies (Table S1) at 4°C overnight, and subsequently with fluorophore conjugated secondary antibodies at room temperature for 1 hour. When fluorophore conjugated primary antibodies (Table S1) were used, the sections were incubated with the antibodies at room temperature for 2 hours. Nuclei were stained with DAPI (Thermo, R37606) at room temperature for 10 min. In between each steps, sections were washed twice with 0.05% Tween-20 in PBS and once with PBS for 5 min for each wash. After the stainings, tumor sections were mounted with mounting medium (Thermo, P36961). For the staining of the thick (30 μm) non-permeabilized section, TritonX-100 was not included. Images were acquired on Leica SP8 confocal microscope or ZEISS Observer microscope and analyzed with ImageJ software.

#### RNA FISH

RNA FISH was done with fresh frozen tumor samples according to user manual of RNAScope Fluorescent Multiplex Reagent kit (ACDBio, 320850). RNAScope probes for mouse *Cxcl12* (422711), *Pdgfra* (480661-C2), and *Krt19* (402941-C3) were used in this study. Images were acquired on Leica SP8 confocal microscope and analyzed with ImageJ.

#### Flow Cytometry

Freshly resected tumors were minced and digested in complete DMEM cell culture medium containing 250 μg/ml Collagenase D (Sigma, 11088858001), 50 μg/ml Liberase DL (Sigma, 05466202001) and 20 μg/ml DNase I (Sigma, 10104159001) at 37°C for 45 min, shaking.

Following lysis of red blood cells in RBC lysis buffer (155 mM NH_4_Cl, 12 mM NaHCO_3_ and 0.1 mM EDTA), cells were passed through 70 μm cell strainer and re-suspend in DMEM medium containing 50 μg/ml DNase I to digest released DNA from RBC cells. After incubation at room temperature for 15 min, the cells were washed and re-suspended in FACS buffer (PBS, 2% FBS and 20 mM HEPES, pH 7.4). For staining, the cells were firstly blocked with Rat anti-mouse CD16/CD32 antibody (BioLegend, 101302) in FACS buffer, shaking at 4°C for 30 min. Then, fluorophore conjugated antibodies (Table S1) were added and incubated for another 30 min in the dark, followed by two washes with FACS buffer. Calcein violet 450 AM (Thermo, 65-0854-39) or DAPI (Thermo, R37606) was used for live cell gating. Data were acquired using a BD LSRForsseta cell analyzer equipped with Diva software 6.0 and were further analyzed with FlowJo.

#### qPCR

Tumor samples of about 30 mg were flash-frozen in liquid nitrogen and stored in −80°C. Total RNA was extracted with RNeasy mini kit (Qiagen, 74106) according to the manual. To make the cDNA library, 1 μg total RNA was reverse transcribed with Taqman Reverse Transcription Kit (Thermo, N8080234), using oligo d(T)s. The cDNA library was then diluted in 100 μl nuclease free water. For qPCR, 5 μl diluted cDNA library was mixed in 384-well plate with specific Taqman probes (Table S2) and Taqman Universal Master Mix II (Thermo, 4440040) to make 10 μl reaction volumes. The qPCRs were run in QuantStudio 6. All the gene expression values were normalized to *Tbp*.

#### RNA-Seq

Subcutaneous tumors were harvest two weeks post cancer cell inoculation. The sequencing libraries were prepared according to Illumina TruSeq Stranded Total RNA Sample Preparation Guide, with certain modifications. Briefly, 4 μg total RNA of each sample was used to make the cDNA library. To enrich DNA fragments at the final step, 12 PCR cycles, instead of 15, was adopted. For library pooling, each cDNA library was diluted to 10 nM and 30 μl of each sample was pooled. The final pooled library was submitted to CSHL Sequencing Core for high-out next generation sequencing, with the sequencing type of single read 76 bp plus bar code.

**Figure S1:**
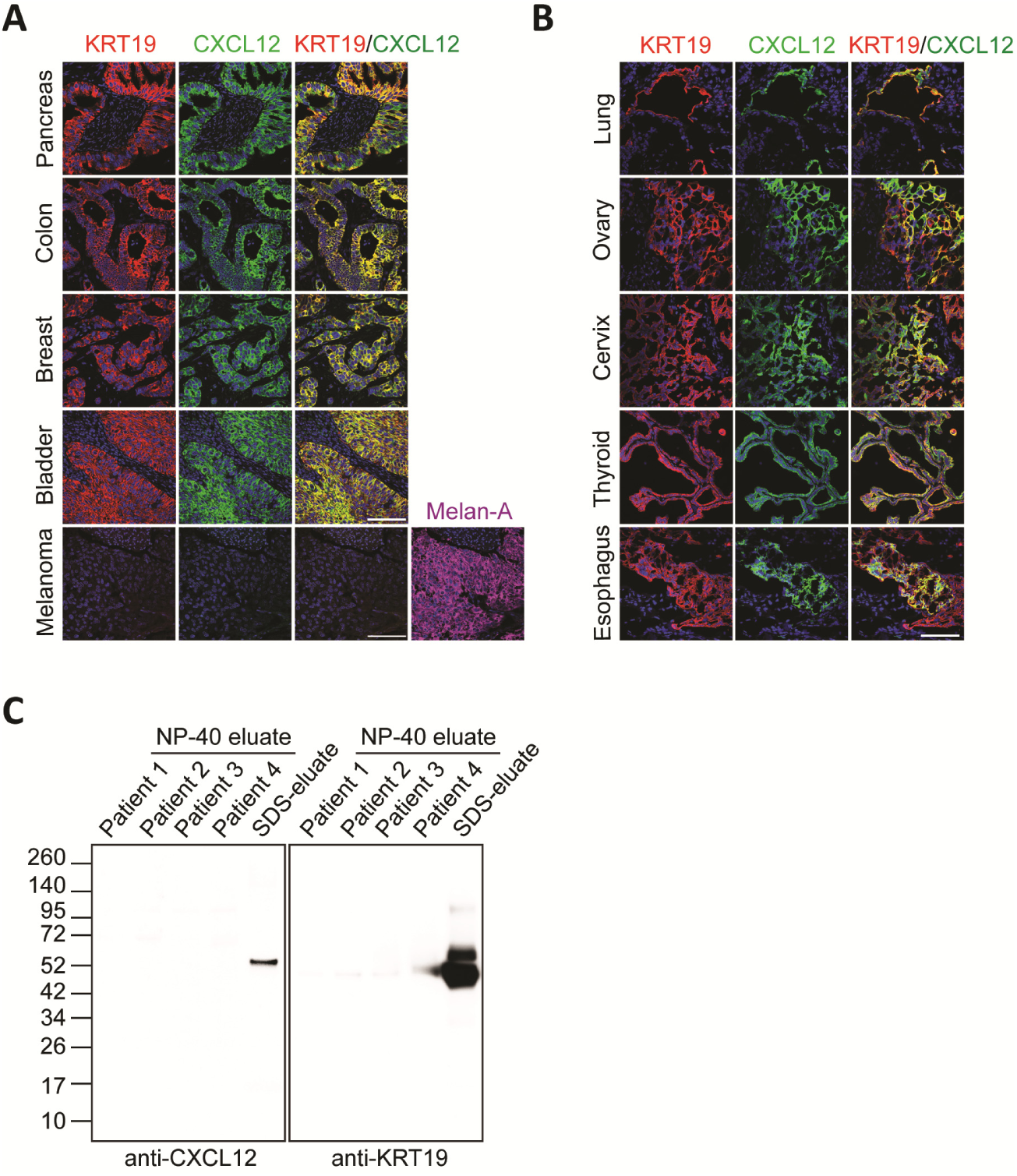
The characterization of the CXCL12-KRT19 coating of human adenocarcinomas. (**A**-**B**) Sections of FFPE human carcinomas and melanoma (A) and frozen human carcinomas (B) were stained with fluorescent antibodies to KRT19 and CXCL12. Scale bars, 100 μm. (**C**) The NP-40 eluates of the four human tumors (Fig. 1A) were subjected to SDS-PAGE and immunoblotting with anti-CXCL12 or anti-KRT19 antibodies. The SDS eluate of patient 2 was applied as a positive control.

**Figure S2:**
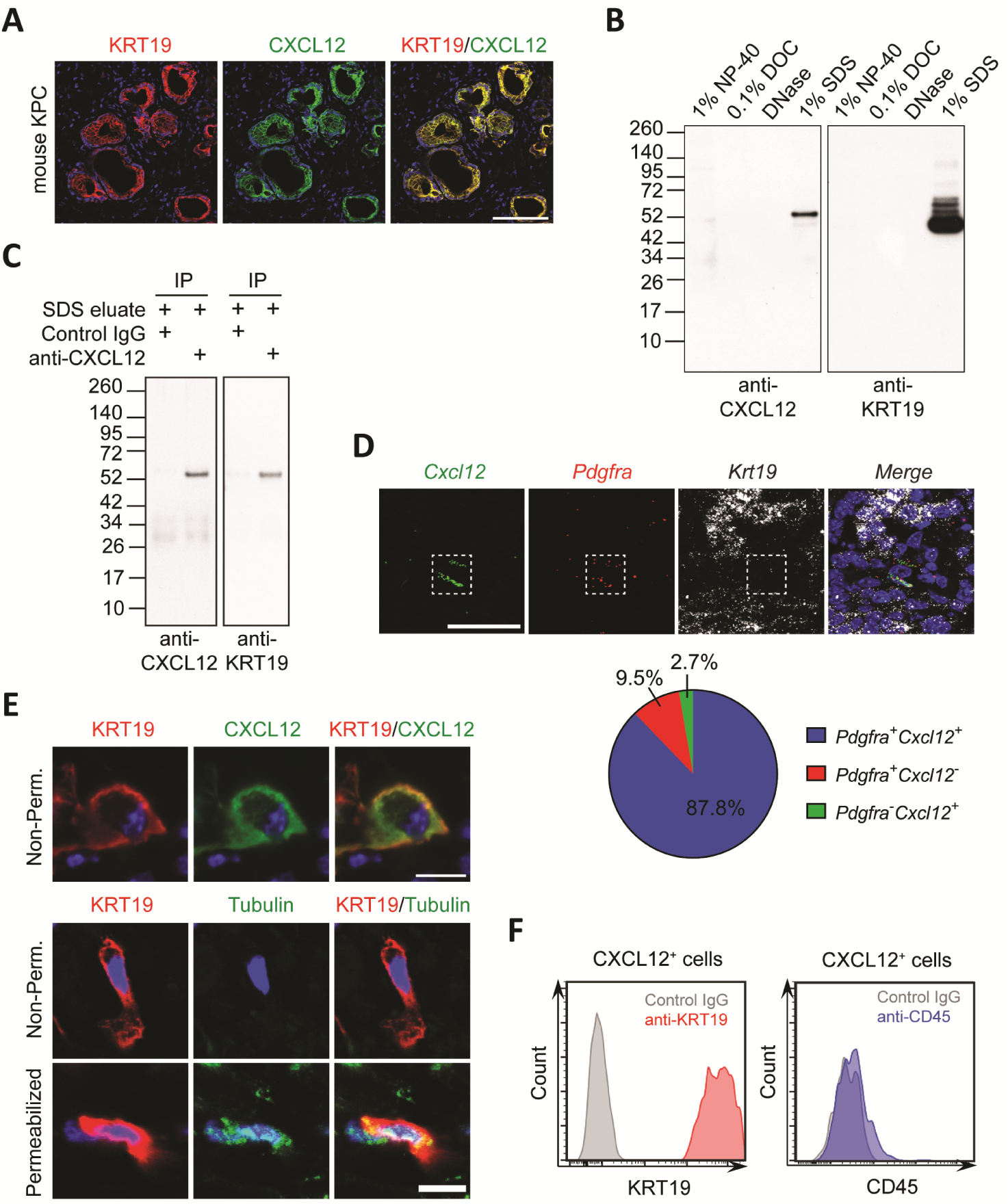
Characterization of the CXCL12-KRT19 coating of mouse PDA tumors. (**A**) A frozen section of mouse autochthonous KPC tumor was stained with fluorescent antibodies to KRT19 and CXCL12. Scale bar, 100 μm. (**B**) Sequential detergent eluates of a mouse subcutaneous PDA were subjected to SDS-PAGE and immunoblotting with anti-KRT19 or anti-CXCL12 antibody. (**C**) The proteins that were immunoprecipitated by control IgG or anti-CXCL12 antibody from the SDS-eluate of a mouse subcutaneous PDA were subjected to SDS-PAGE and immunoblotting with anti-CXCL12 or anti-KRT19 antibodies. (**D**) The mRNAs of *Cxcl12, Pdgfra* (indicating CAFs) and *Krt19* (PDA cells) were detected by RNA fluorescent in situ hybridization (FISH) in sections of a mouse subcutaneous PDA. The pie chart shows relative proportions of the cells that are *Pdgfra*^+^, or *Cxcl12*^+^ or double positive. Scale bar, 50 μm. (**E**) 30 μm-thick sections of a mouse subcutaneous PDA were permeabilized with Triton X-100, or not, were stained with fluorescent antibodies to KRT19, CXCL12, and tubulin. Scale bars, 10 μm. (**F**) Intact, DAPI-excluding dissociated cells from a mouse subcutaneous PDA tumor that had been stained with fluorescent antibodies to CXCL12, KRT19 and CD45, were assessed by flow cytometry.

**Figure S3:**
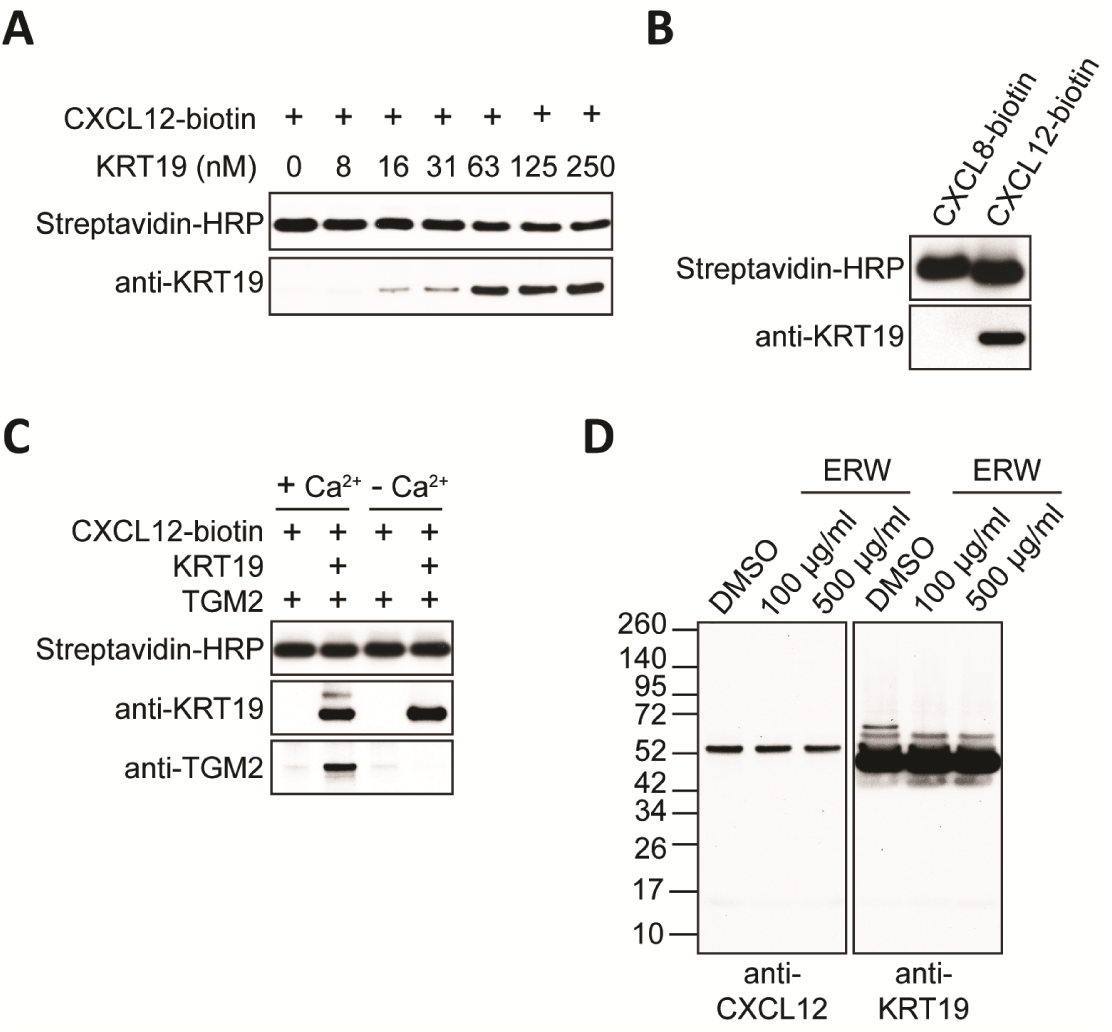
The interactions among CXCL12, KRT19, and TGM2. (**A**) In the experiment depicted in Fig. 1D, KRT19 bound to CXCL12 was measured by immunoblot analysis of the eluates of streptavidin beads. (**B**) Biotinylated human CXCL8 or CXCL12 that was immobilized on streptavidin beads was incubated with KRT19, and bound proteins were subjected to SDS-PAGE and blotting with streptavidin-HRP or anti-KRT19 antibody. (**C**) CXCL12-biotin or the preformed CXCL12-biotin/KRT19 complex that was immobilized on streptavidin beads was incubated with TGM2, and bound proteins were subjected to SDS-PAGE and blotting with streptavidin-HRP, anti-KRT19, and anti-TGM2, as indicated. (**D**) Sequential extractions of mouse subcutaneous PDA were performed in the presence of 100 μg/ml, 500 μg/ml ERW1041E, or DMSO, and the SDS eluates were analyzed by SDS-PAGE and immunoblotting with anti-CXCL12 or anti-KRT19 antibodies.

**Figure S4:**
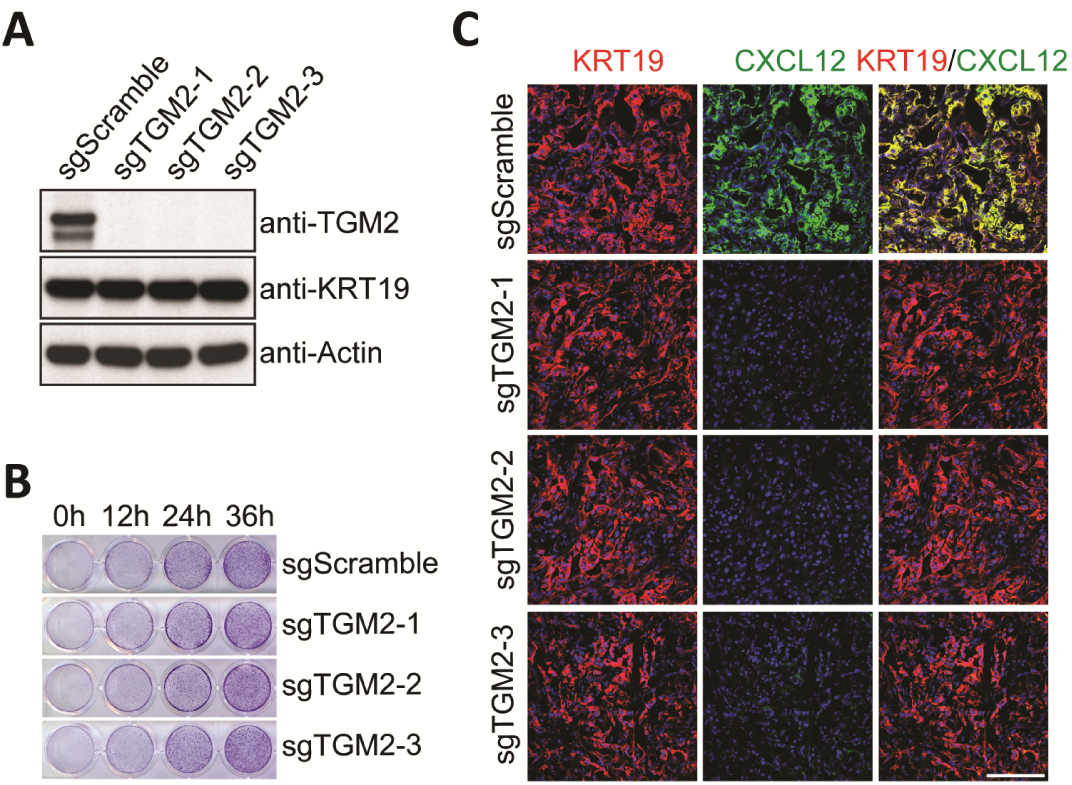
*Tgm2*-edited PDA cells and the formation of the CXCL12-KRT19 coating. (**A**) Lysates from mouse PDA cell lines that were CRISPR/Cas9 edited with scramble sgRNA or three different *Tgm2* sgRNAs were subjected to SDS-PAGE and immunoblotting with anti-TGM2, anti-KRT19 or anti-actin antibodies. (**B**) *In vitro* growth of sgScramble and sgTGM2-edited PDA cells was assessed by staining every 12 hours with crystal violet. (**C**) Tissue sections of subcutaneous tumors formed by control and the three *Tgm2*-edited PDA cell lines were stained with fluorescent antibodies to KRT19 and CXCL12, respectively. Scale bar, 100 μm.

**Figure S5:**
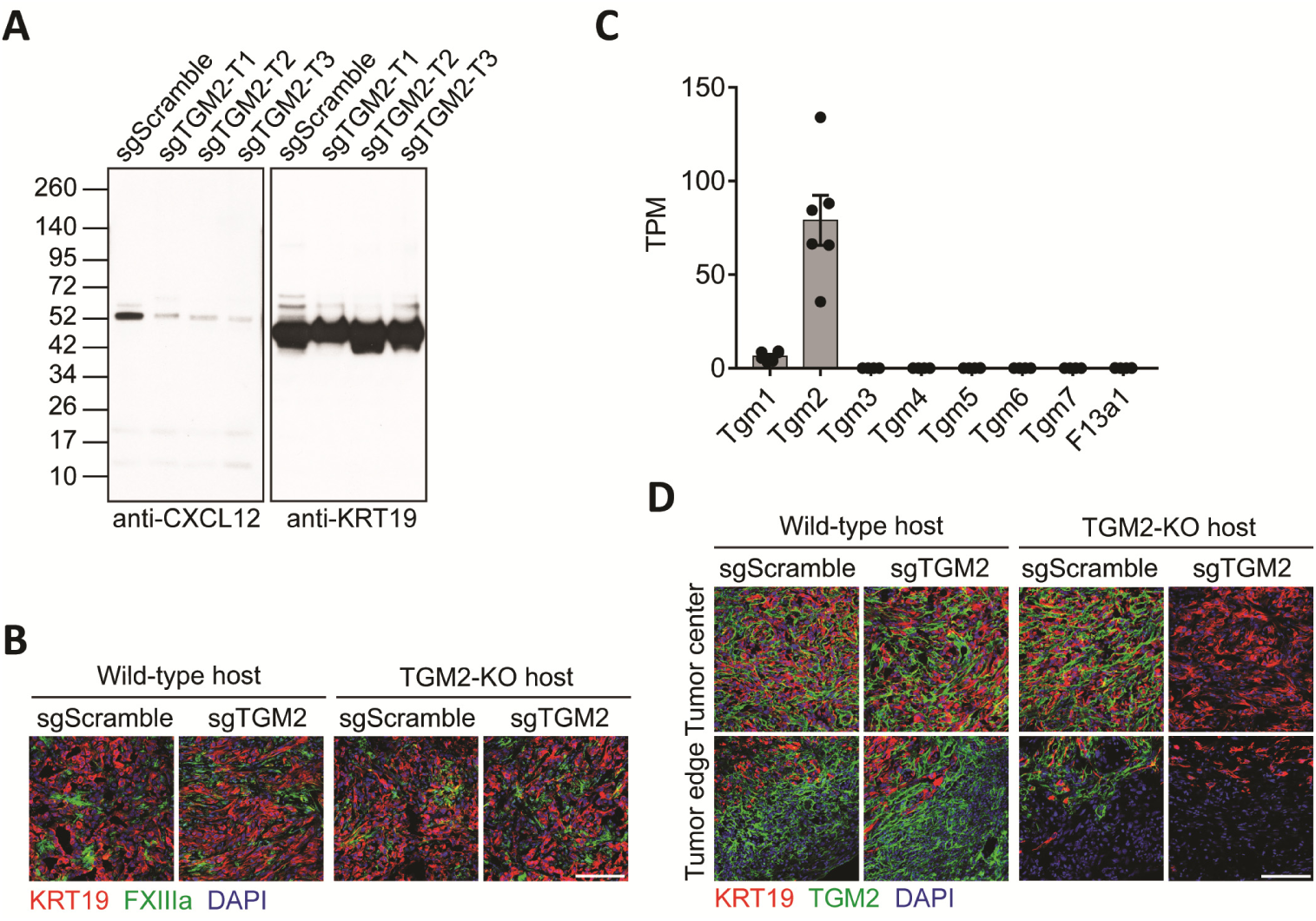
The role of TGM2 expression by mouse PDA cells in the formation of the CXCL12-KRT19 coating. (**A**) SDS eluates of subcutaneous tumors formed with sgScramble or sgTGM2-edited PDA cells in wild-type hosts were subjected to SDS-PAGE and immunoblotting with anti-CXCL12 or anti-KRT19 antibodies. (**B**) Sections of subcutaneous tumors formed with sgScramble or sgTGM2-edited PDA cell lines, in wild-type hosts or TGM2-KO hosts, were stained with fluorescent anti-KRT19 and anti-FXIIIa antibodies. (**C**) mRNA levels of members of the transglutaminase family in mouse PDA cancer cell organoids are shown (*11*). (**D**) Subcutaneous tumors formed with sgScramble or sgTGM2-edited PDA cell lines, in wild-type hosts or TGM2-KO hosts, were stained with fluorescent anti-KRT19 and anti-TGM2 antibodies. Scale bars, 100 µm.

**Figure S6:**
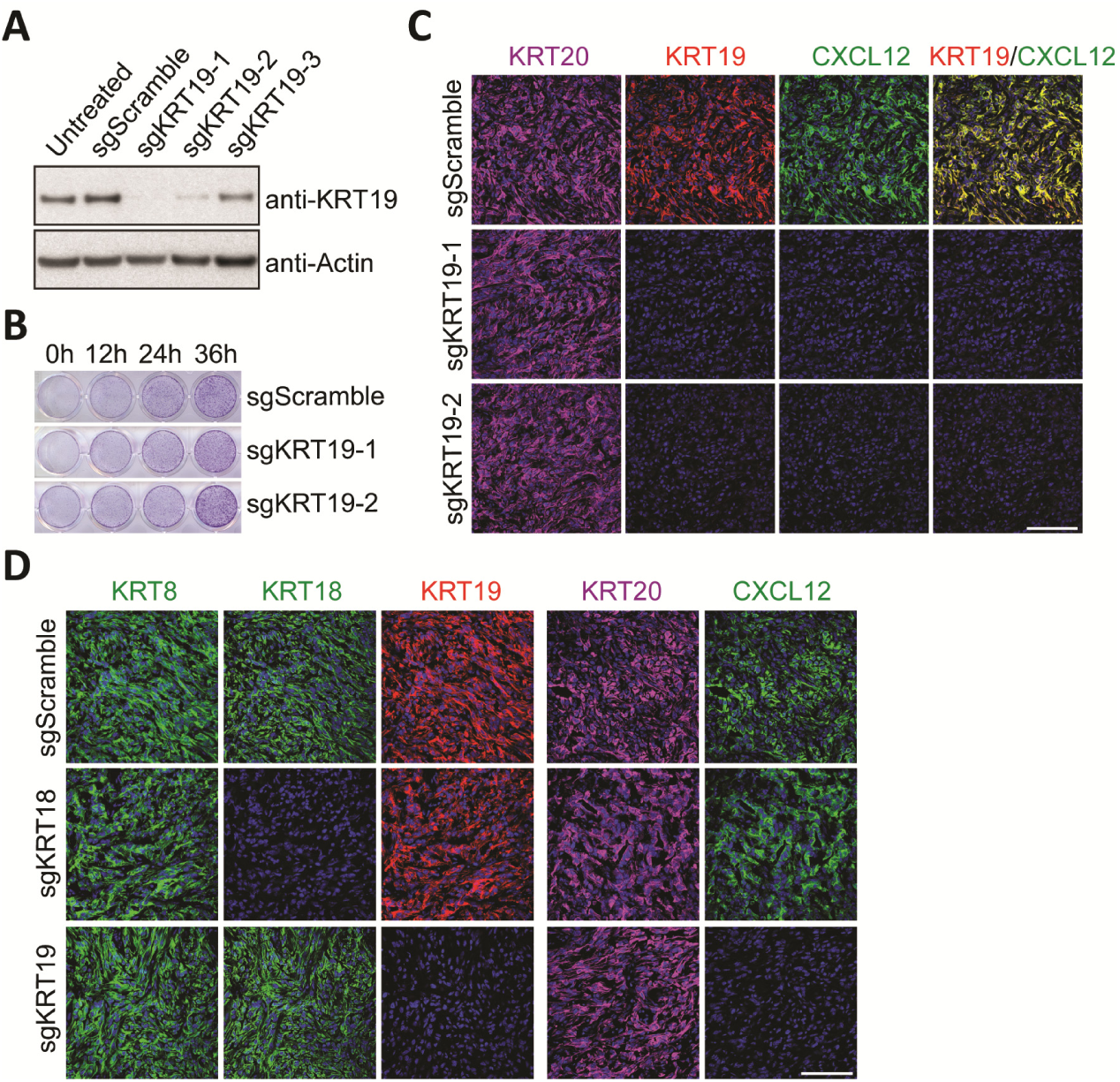
*Krt19*-edited PDA cells and the formation of the CXCL12-KRT19 coating. (**A**) Lysates from mouse PDA cell lines that were CRISPR/Cas9 edited with scramble sgRNA or three different *Krt19* sgRNAs were subjected to SDS-PAGE and immunoblotting with anti-KRT19 or anti-actin antibodies. (**B**) Crystal violet stains measuring the *in vitro* growth of sgScramble and sgKRT19-edited PDA cells are shown. (**C**) Tissue sections of subcutaneous tumors formed by control and the three *Krt19*-edited PDA cell lines were stained with fluorescent antibodies to KRT20 to reveal cancer cells, KRT19 and CXCL12, respectively. (**D**) Sections of subcutaneous tumors formed with sgScramble-, sgKRT18- or sgKRT19-edited PDA cells were stained with fluorescent antibodies to KRT8, KRT18, KRT19, KRT20 and CXCL12, respectively. Scale bars, 100 μm.

**Figure S7:**
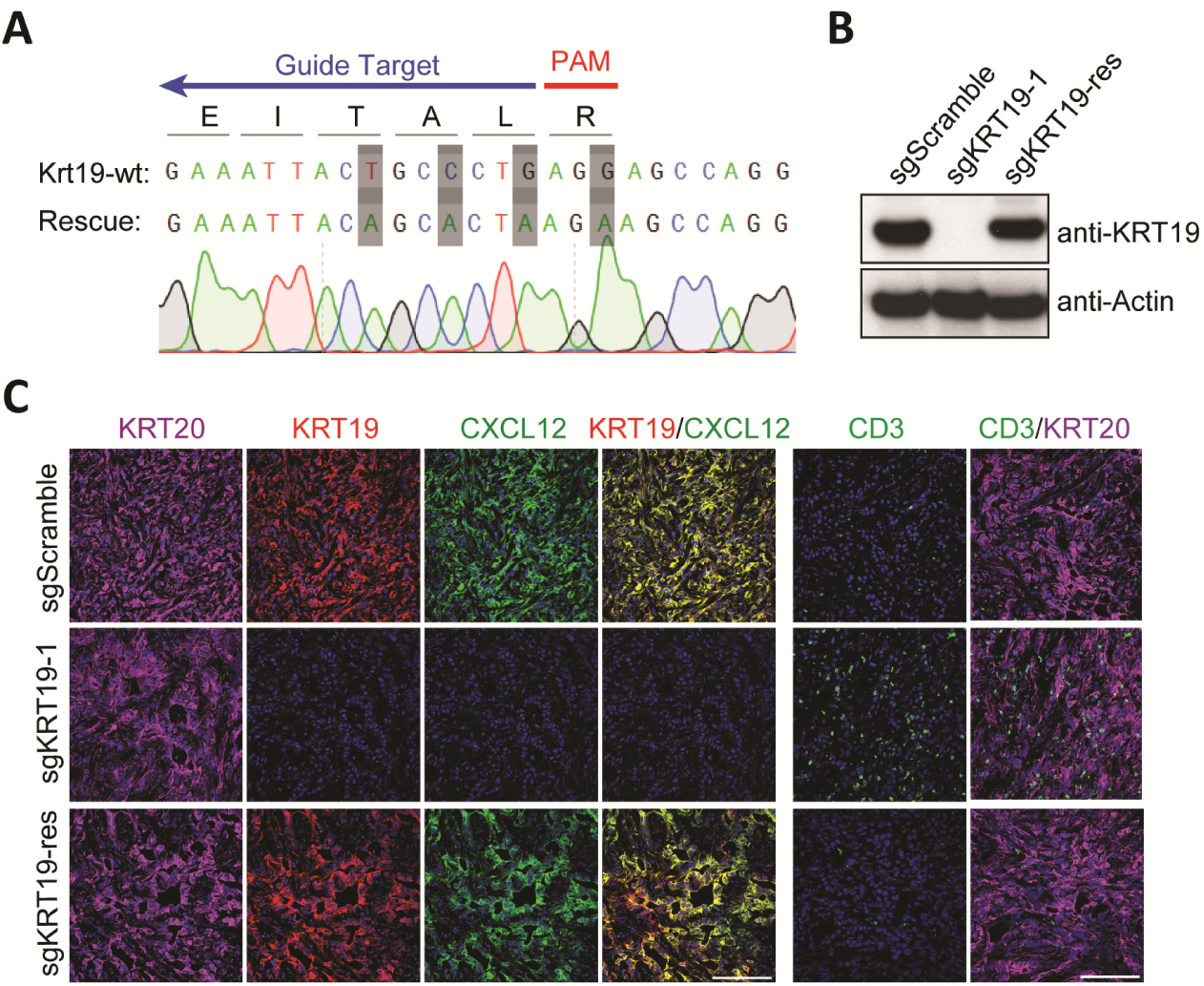
Rescue of KRT19 expression in *Krt19*-edited PDA cells. (**A**) Sequencing of the KRT19 rescue construct where four synonymous mutations, shown in black boxes, were introduced to prevent the binding of sgKRT19-1. (**B**) PDA cell lysates were subjected to SDS-PAGE and immunoblotting with anti-KRT19 and anti-actin antibodies. (**C**) Tissue sections of subcutaneous tumors formed with the sgScramble PDA cells, sgKRT19-edited PDA cells, and the KRT19-rescue PDA cells were stained with fluorescent antibodies to KRT20 to reveal cancer cells, KRT19, CXCL12, and CD3, respectively. Scale bars, 100 μm.

**Figure S8:**
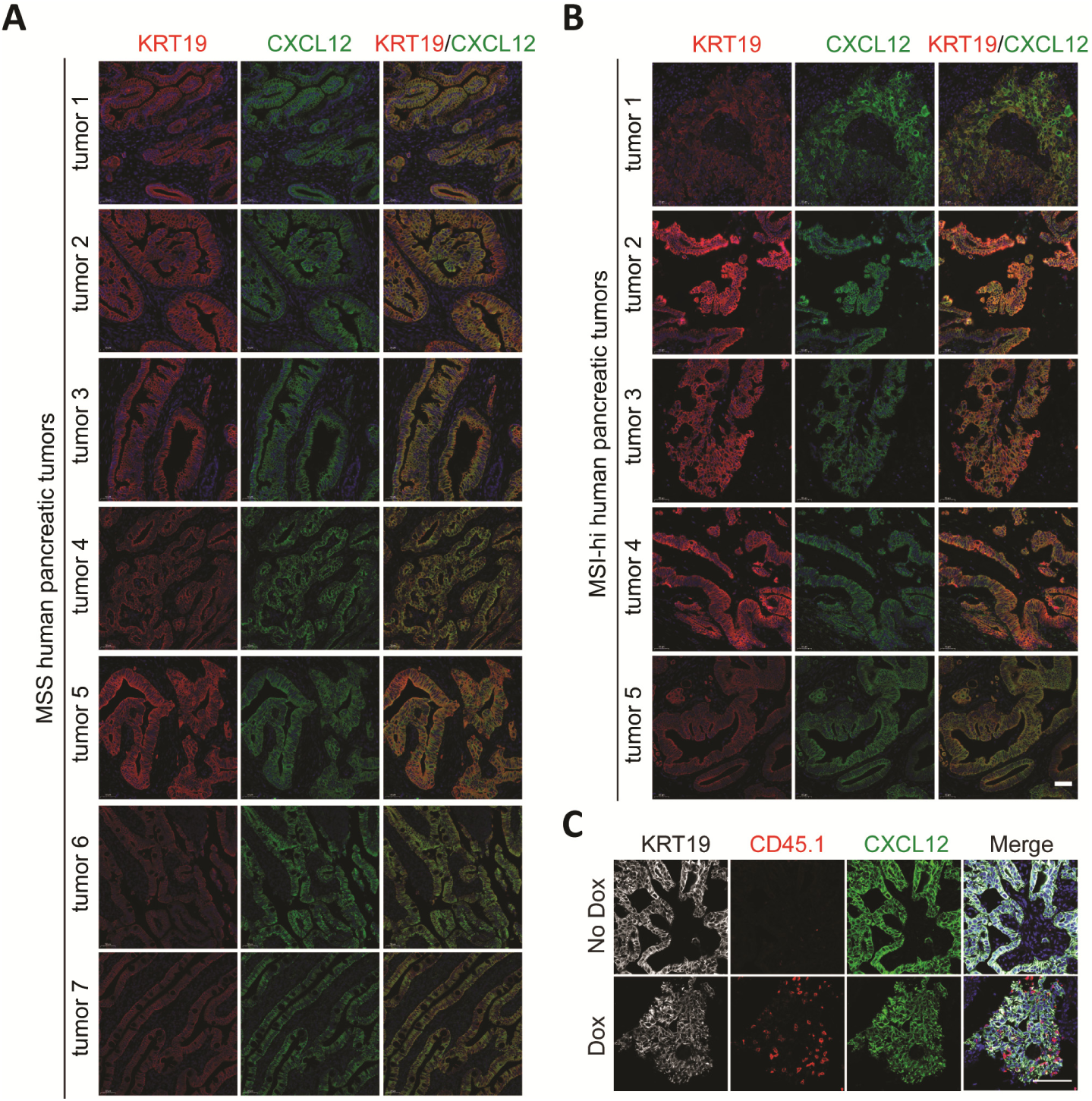
The CXCL12-KRT19 coating of cancer cells in MSS and MSI-Hi PDA and in mouse hepatic PDA metastases. (**A**-**B**) Sections of FFPE human MSS and MSI-Hi PDA were stained with fluorescent antibodies to KRT19 and CXCL12. (**C**) Sections of mouse hepatic PDA metastases formed without and with Dox-induced OVA expression were stained with fluorescent antibodies to KRT19, CXCL12, and CD45.1. The CD45.1^+^ OT-I cells were adoptively transferred to mice before the intra-portal injection of the PDA cells. Scale bars, 100 μm.

**Figure S9:**
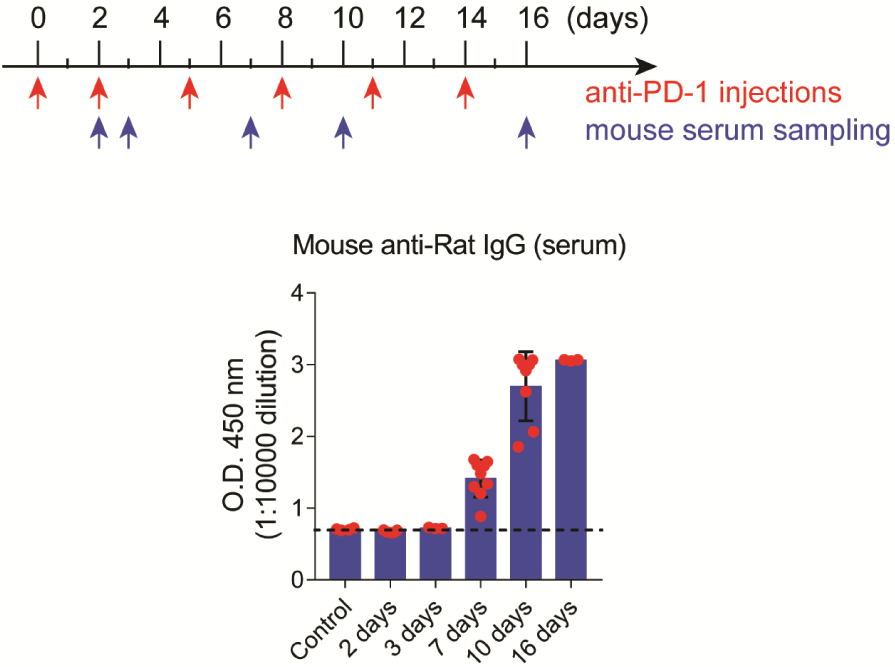
Development of neutralizing anti-rat antibodies in mice receiving rat anti-mouse PD-1 antibody. Mouse anti-rat IgG in blood serum of mice receiving rat anti-mouse PD-1 antibody was measured by means of ELISA. 200 μg rat anti-mouse PD-1 antibody was administered by intra-peritoneal injections at indicated days (red arrows).

**Figure S10:**
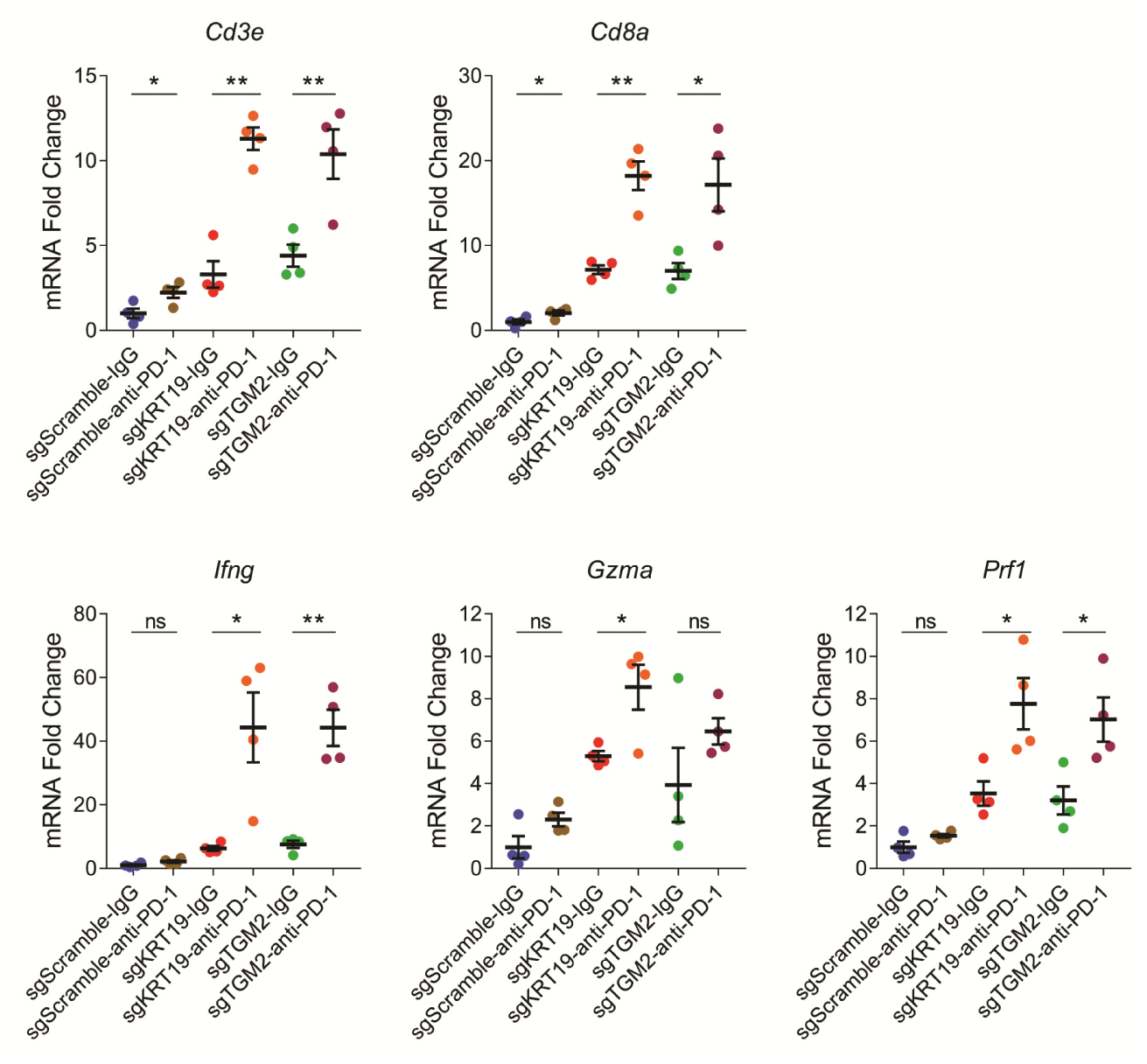
Gene expression analysis of the subcutaneous PDA tumors treated with anti-PD-1 antibody. PDA-bearing mice were treated with 200 μg isotype IgG or anti-PD-1 antibody at day 13 and day 15 after cancer cell inoculation. Mice were sacrificed at day17, and samples of the tumors were obtained. The mRNA levels of each gene in the tumor samples were measured by qPCR. Expression of each gene in sgScramble tumors from mice that had been treated with isotype IgG were normalized to 1. Mean ± SEM; * *p*<0.05, ** *p*<0.01, Student’s *t* test.

**Figure S11:**
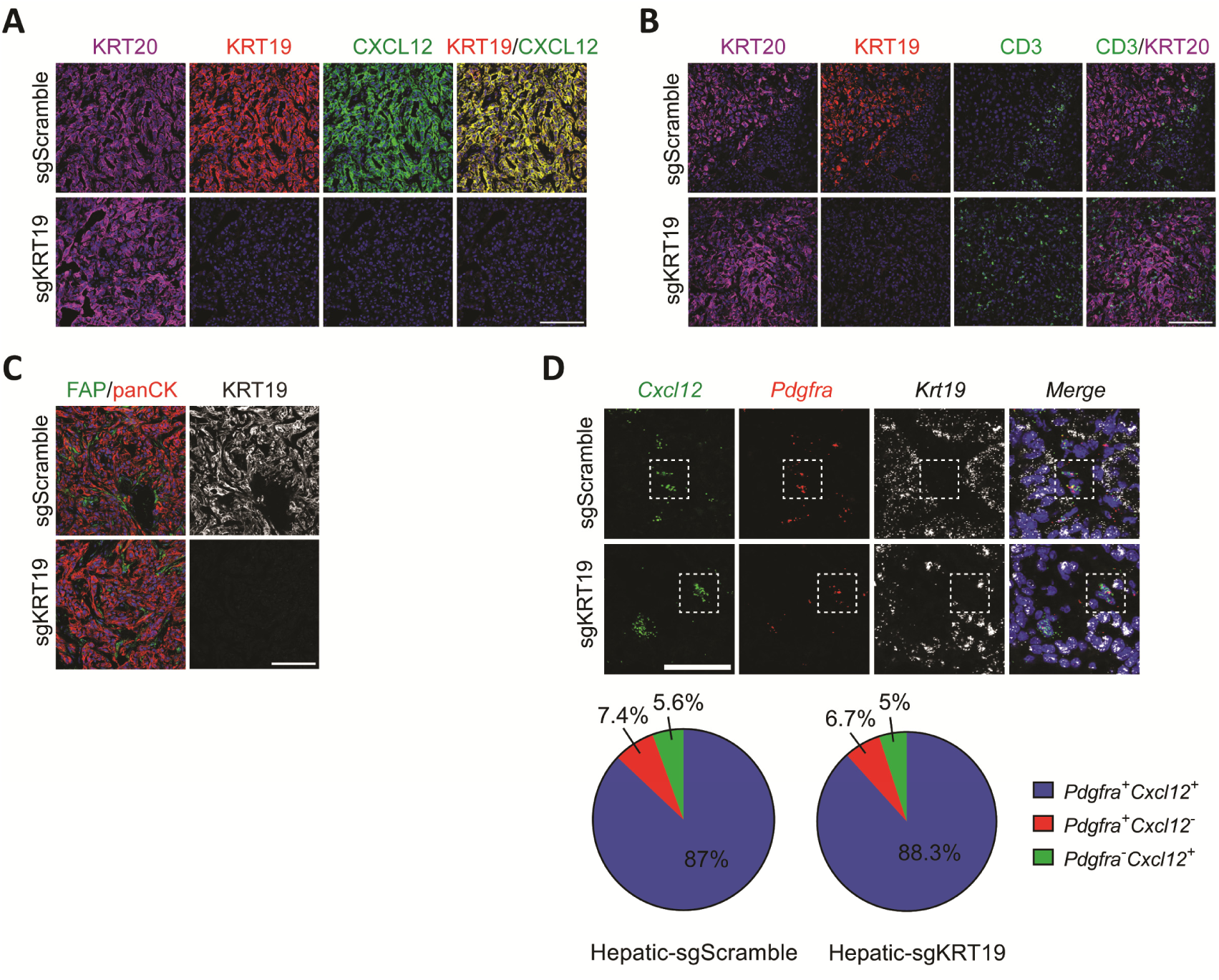
Characterization of *Krt19*-edited mouse PDA hepatic metastases. (**A**-**C**) Sections of the control and *Krt19*-edited PDA metastases were stained with fluorescent antibodies to KRT20, KRT19, CXCL12, CD3, fibroblast activation protein (FAP), and pan-cytokertain (CK), respectively. Scale bars, 100 μm. (**D**) mRNAs for *Cxcl12, Pdgfra* and *Krt19* were detected by RNA FISH in sections of hepatic metastases. The pie chart shows the relative proportions of the cells that are *Pdgfra*^+^, or *Cxcl12*^+^ or double positive. Scale bar, 50 μm.

**Figure S12:**
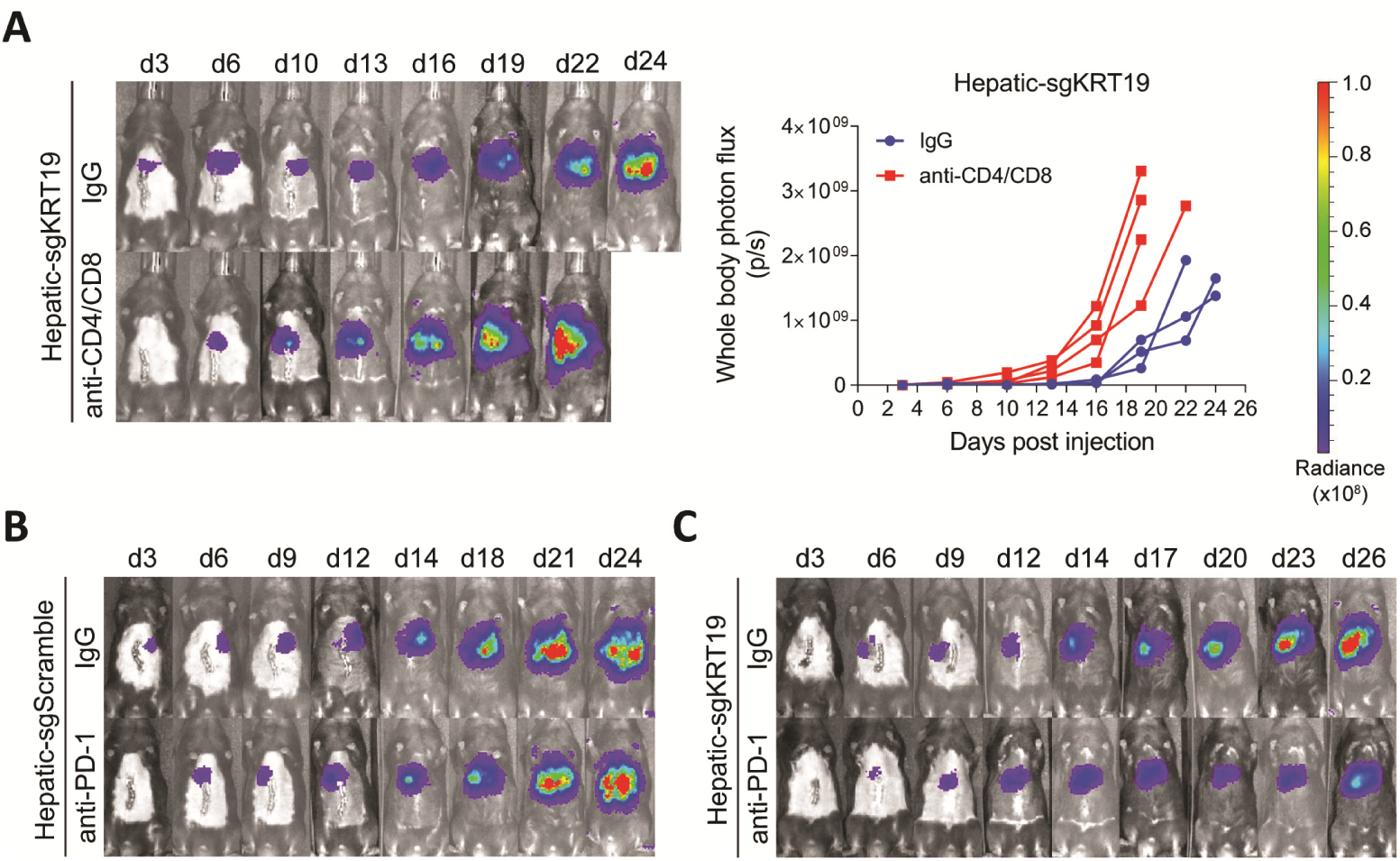
Bioluminescent images of hepatic metastases formed with luciferase-expressing control and *Krt19*-edited PDA cells. (**A**) Hepatic metastases were formed by injecting 2.5×10^4^ *Krt19*-edited PDA cells into the portal vein at day 0 in mice that had been pre-treated with non-immune IgG or depleting antibodies to CD4 and CD8, and growth of the metastases was measured by bioluminescent imaging. (**B**-**C**) Representative bioluminescence images of the mice in which metastasis growth was depicted in Fig. 4B.

**Figure S13:**
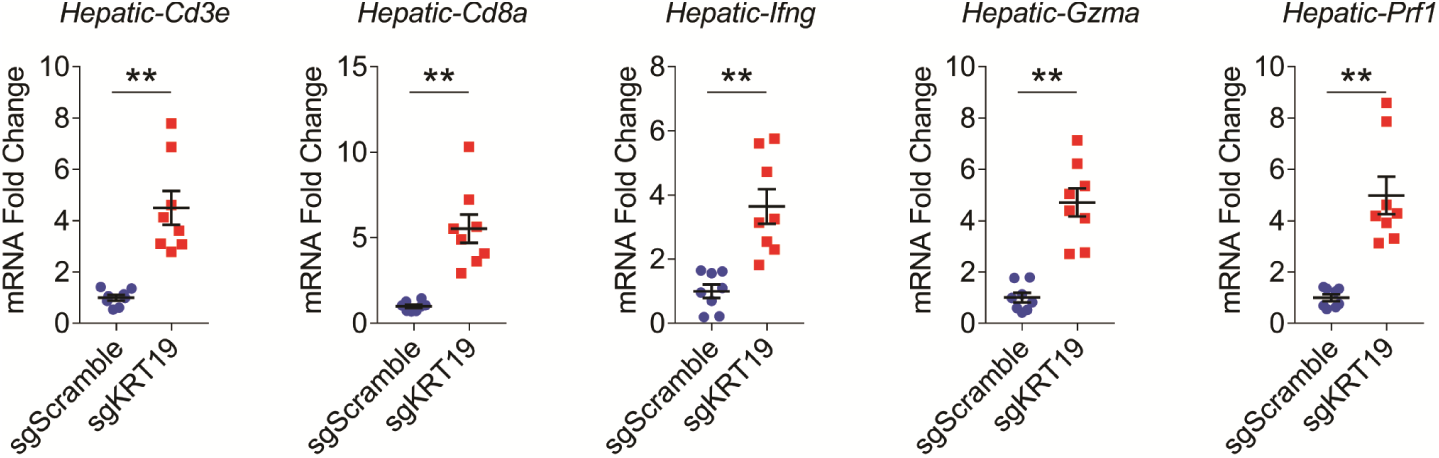
Expression of immune genes in hepatic metastases formed with control (sgScramble) and *Krt19*-edited (sgKRT19) PDA cells. The mRNA levels were measured by qPCR and expression of each gene was normalized to that of the control levels, which were assigned a value of 1. Mean ± SEM; ** *p*<0.01, Student’s *t* test.

**Table S1:** Primary antibody list.

| **Antibody** | **Vendor** | **Catalog NO.** | **Reactivity** | **Application** |
| --- | --- | --- | --- | --- |
| KRT19-AF568 | Abcam | ab203445 | Hs, Mm | IF, FC |
| KRT19-AF647 | Abcam | ab192980 | Hs, Mm | IF, FC |
| KRT8-AF647 | Abcam | ab192468 | Mm | IF |
| CXCL12-FITC | R&D | IC350F | Hs, Mm | IF (Frozen), FC |
| Melan-A-AF647 | NOVUS | NBP2-46603AF647 | Hs | IF |
| CD45-BV510 | BioLegend | 103138 | Mm | FC |
| CD3-AF647 | BioLegend | 100209 | Mm | IF |
| Tubulin-AF488 | Cell Signaling | 36234S | Mm | IF |
| panCK-AF647 | NOVUS | NBP1-48348AF647 | Mm | IF |
| Mm-anti-CXCL12 | R&D | MAB350 | Hs, Mm | IF (FFPE) |
| Rb-anti-CXCL12 | Peprotech | 500-P87A | Hs, Mm | IF (FFPE), WB |
| Rb-anti-CXCL12 | LSBio | LS-C48888 | Hs, Mm | IP |
| Rb-anti-KRT19 | Abcam | ab76539 | Hs | IP |
| Rb-anti-KRT19 | Abcam | ab52625 | Hs, Mm | IF, WB |
| Rt-anti-KRT19 | DHSB | TROMA-III | Mm | WB |
| Rb-anti-KRT20 | Abcam | ab230524 | Mm | IF |
| Mm-anti-KRT18 | Abcam | ab668 | Mm | IF, WB |
| Rb-anti-KRT8 | Abcam | ab53280 | Mm | IF, WB |
| Sh-anti-TGM2 | R&D | AF5418 | Mm | IF, WB |
| Mm-anti-FXIIIa | Abcam | ab1834 | Mm | IF |
| Rb-anti-Actin | Abcam | ab8227 | Mm | WB |
| Rb-anti-FAP | Millipore | ABT11 | Mm | IF |
| Mm-anti-CD45.1 | BioLegend | 110702 | Mm | IF |
| Rt-anti-CD3 | BioLegend | 100202 | Mm | IF |
| Rt-anti-CD8 | BioXcell | BP0061 | Mm | T cell Depletion |
| Rt-anti-CD4 | BioXcell | BP0003 | Mm | T cell Depletion |
| Rt-anti-PD-1 | BioXcell | BP0273 | Mm | PD-1 blocking |

**Table S2:**
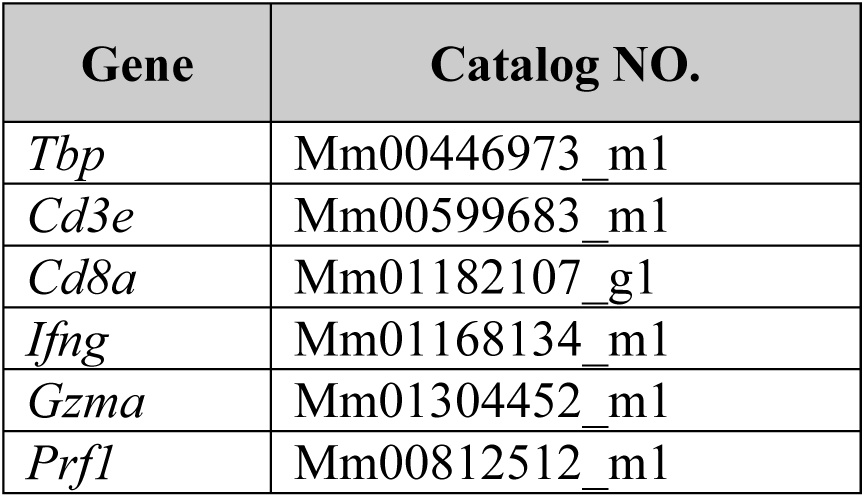
Taqman probes for qPCR (Thermo)

## References and Notes

1. J. A. Joyce, D. T. Fearon, T cell exclusion, immune privilege, and the tumor microenvironment. Science 348, 74–80 (2015).

2. C. Feig et al., Targeting CXCL12 from FAP-expressing carcinoma-associated fibroblasts synergizes with anti-PD-L1 immunotherapy in pancreatic cancer. Proc Natl Acad Sci U S A 110, 20212–20217 (2013).

3. M. S. Daniele Biasci, Claire Connell, Zhikai Wang, Ya Gao, James Thaventhiran, Bristi Basu, Lukasz Magiera, Isaac Johnson, Lisa Bax, Aarthi Gopinathan, Christopher Isherwood, Ferdia Gallagher, Maria Pawula, Irena Hudecova, Davina Gale, Nitzan Rosenfeld, Petros Barmpounakis, Elizabeta Popa, Rebecca Brais, Edmund Godfrey, Fraz Mir, Frances Richards, Douglas Fearon, CXCR4 inhibition in human pancreatic and colorectal cancers induces an integrated immune response. medRxixv doi: https://doi.org/10.1101/2020.07.08.20129361, (2020).

4. B. Bockorny et al., BL-8040, a CXCR4 antagonist, in combination with pembrolizumab and chemotherapy for pancreatic cancer: the COMBAT trial. Nat Med 26, 878–885 (2020).

5. C. Alix-Panabieres et al., Full-length cytokeratin-19 is released by human tumor cells: a potential role in metastatic progression of breast cancer. Breast Cancer Res 11, R39 (2009).

6. C. A. Iacobuzio-Donahue et al., Exploration of global gene expression patterns in pancreatic adenocarcinoma using cDNA microarrays. Am J Pathol 162, 1151–1162 (2003).

7. O. Ayinde, Z. Wang, M. Griffin, Tissue transglutaminase induces EpithelialMesenchymal-Transition and the acquisition of stem cell like characteristics in colorectal cancer cells. Oncotarget 8, 20025–20041 (2017).

8. L. Dafik, M. Albertelli, J. Stamnaes, L. M. Sollid, C. Khosla, Activation and inhibition of transglutaminase 2 in mice. PLoS One 7, e30642 (2012).

9. D. M. Pinkas, P. Strop, A. T. Brunger, C. Khosla, Transglutaminase 2 undergoes a large conformational change upon activation. PLoS Biol 5, e327 (2007).

10. N. Nanda et al., Targeted inactivation of Gh/tissue transglutaminase II. J Biol Chem 276, 20673–20678 (2001).

11. S. F. Boj et al., Organoid models of human and mouse ductal pancreatic cancer. Cell 160, 324–338 (2015).

12. D. T. Le et al., Mismatch repair deficiency predicts response of solid tumors to PD-1 blockade. Science 357, 409–413 (2017).

## References

(13) K. Suzuki et al., In vivo genome editing via CRISPR/Cas9 mediated homology-independent targeted integration. Nature 540, 144–149 (2016).

(14) S. R. Hingorani et al., Trp53R172H and KrasG12D cooperate to promote chromosomal instability and widely metastatic pancreatic ductal adenocarcinoma in mice. Cancer Cell 7, 469–483 (2005).

